# Multiple Flagellin Proteins Have Distinct and Synergistic Roles in *Agrobacterium tumefaciens* Motility

**DOI:** 10.1101/335265

**Authors:** Bitan Mohari, Melene A. Thompson, Jonathan C Trinidad, Clay Fuqua

## Abstract

Rotary flagella propel bacteria through liquid and across semi-solid environments. Flagella are composed of the basal body that constitutes the motor for rotation, the curved hook that connects to the basal body, and the flagellar filament that propels the cell. Flagellar filaments can be comprised of a single flagellin protein such as in *Escherichia coli* or with multiple flagellins such is in *Agrobacterium tumefaciens*. The four distinct flagellins FlaA, FlaB, FlaC and FlaD produced by wild type *A. tumefaciens*, are not redundant in function, but have specific properties. FlaA and FlaB are much more abundant than FlaC and FlaD and are readily observable in mature flagellar filaments, when either FlaA or FlaB is fluorescently labeled. Cells having FlaA with any one of the other three flagellins can generate functional filaments and thus are motile, but FlaA alone cannot constitute a functional filament. In *flaA* mutants that manifest swimming deficiencies, there are multiple ways by which these mutations can be phenotypically suppressed. These suppressor mutations primarily occur within or upstream of the *flaB* flagellin gene or in the transcriptional factor *sciP* regulating flagellar expression. The helical conformation of the flagellar filament appears to require a key asparagine residue present in FlaA and absent in other flagellins. However, FlaB can be spontaneously mutated to render helical flagella in absence of FlaA, reflecting their overall similarity and perhaps the subtle differences in the specific functions they have evolved to fulfill.

**Importance:** Flagellins are abundant bacterial proteins comprising the flagellar filaments that propel bacterial movement. Several members of the Alphaproteobacterial group express multiple flagellins, in contrast to model systems such as *Escherichia coli* that has only one flagellin protein. The plant pathogen *Agrobacterium tumefaciens* has four flagellins, the abundant and readily detected FlaA and FlaB, and lower levels of FlaC and FlaD. Mutational analysis reveals that FlaA requires at least one of the other flagellins to function - *flaA* mutants produce non-helical flagella and cannot swim efficiently. Suppressor mutations can rescue this swimming defect through mutations in the remaining flagellins, including structural changes imparting flagellar helical shape, and putative regulators. Our findings shed light on how multiple flagellins contribute to motility.

## Introduction

Flagella typically function as a means of bacterial locomotion in liquid environments, by a discontinuous pattern of swimming comprised of straight runs and directional changes known as tumbles. Other forms of motility include swarming, which also requires flagella, and gliding, twitching, and sliding which do not. Flagella are extracellular helical filaments with a curved hook serving as a universal joint between the filament and the rotary basal body embedded in the cellular membrane (1). The basal body generates power and rotates the flagellum and the hook is important for rendering the optimum angle for propulsion. Flagellar filaments are composed of up to 30,000 subunits of the protein flagellin (2). For bacteria with multiple flagella, these form a bundle to propel forward movement and unbundle to promote cellular re-orientation (3). Flagellar rotation in enteric bacteria such as *Escherichia coli* and *Salmonella typhimurium* is bidirectional; (counterclockwise promoting bundling and straight swimming, and reversals to clockwise rotation disrupting the flagellar bundle, and causing tumbles). However, in members of the family *Rhizobiaceae*, including *Sinorhizobium meliloti* and *Agrobacterium tumefaciens*, the rotation is unidirectional and always clockwise, with disruption of bundles occurring due to discordance of flagellar rotation rates (4). The helical shape of the flagellar filament is of utmost importance for these dynamic aspects of propulsion (5).

Bacterial filaments can be categorized as plain or complex depending on whether they are made of one or multiple kinds of flagellin protein (6). In well studied systems, such as peritrichously flagellated *E. coli*, there are multiple filaments comprised of the FliC flagellin protein (7, 8). Similarly the alphaproteobacterium *Rhodobacter sphaeroides* has one flagellum (laterally positioned on the cell) that is made up of a single flagellin protein (9, 10). On the other hand, examples of bacteria with filaments made of multiple flagellins include *Caulobacter crescentus*, which has a single polar flagellum, but remarkably six flagellin proteins (11). Within the *Rhizobiaceae* family *Sinorhizobium meliloti, Rhizobium leguminosarum, Agrobacterium* sp. H13-3, and *Agrobacterium tumefaciens* all encode multiple flagellins. *A. tumefaciens* mutated in three of its four flagellin genes is reduced in virulence by about 38% and *Agrobacterium* sp. H13-3 lacking all of its three flagellin genes is resistant to flagella-specific phage infection (12-16). Flagella with simple, single flagellin filaments exhibit structural polymorphisms during the course of normal flagellar propulsion and rotational switching, whereas those with complex filaments do not exhibit these polymorphisms unless they are exposed to extreme conditions of pH and ionic strengths (13, 17). Studies in *C. crescentus* with six different flagellin genes suggest that multiple flagellins can have a certain level of redundancy (11). In some cases there can be a single predominant flagellin required for flagellar function, such as those for *S. meliloti, Agrobacterium* sp. H13-3 and *Rhizobium leguminosarum* (12, 14). Multiple flagellins can also be differentially regulated as in spirochetes, *Vibrio cholerae*, and *C. crescentus* (11, 18, 19).

*A. tumefaciens* has a lophotrichous arrangement of 5-6 flagellar filaments hypothesized to be comprised of four flagellins. The flagellins all share significant sequence similarity (Table 1; Fig. S1). The functions and coordination of the flagellins are however, not well studied. FlaA, FlaB and FlaC are encoded in a single gene cluster and are of similar molecular mass: (FlaA, 306 aa; FlaB, 320 aa; FlaC, 313 aa). In contrast, FlaD is the most dissimilar, located roughly 15 kb distal to the *flaABC* cluster and is oriented in the opposite direction, relative to the other flagellins (Fig. S2). FlaD is 430 aa, with a large internal segment that is not shared with the other flagellins. FlaA is reported to be the major flagellin, as disruption of the gene severely compromised motility, reportedly resulting in vestigial stubs instead of flagellar filaments (16, 20). Other single flagellin mutants were shown to be attenuated in motility that produced filaments with some structural abnormalities. Mutants were generated either via transposon or antibiotic cassette insertions, and thus are prone to polar effects on downstream genes and partial gene copies, confounding the results reported for these flagellins and making it difficult to evaluate individual gene function (16, 20).

**Table 1.** Comparison of *A. tumefaciens* flagellin genes

| Protein | Gene<sup>a</sup> | Length<br>(aa) | % Sequence Identity (Conservation) with: |  |  |  |
| --- | --- | --- | --- | --- | --- | --- |
|  |  |  | FlaA | FlaB | FlaC | FlaD |
| FlaA | Atu0545 | 306 |  | 62 (76) | 60 (74) | 54 (69) |
| FlaB | Atu0543 | 320 |  |  | 62 (77) | 55 (70) |
| FlaC | Atu0542 | 313 |  |  |  | 57 (74) |
| FlaD | Atu0567 | 430 |  |  |  |  |

<sup>a</sup> Atu numbers as available through BioCyc database (https://biocyc.org/)

*S. meliloti* and *Agrobacterium* sp H13-3 have been studied for their multiple flagellins. *S. meliloti* has four flagellins like *A. tumefaciens*, and *Agrobacterium* sp H13-3 has three flagellins FlaA, FlaB and FlaD (12). Mutational analysis suggested that FlaA along with another secondary flagellin are required to assemble a functional filament and render proficient motility. Although these three taxa are related, amino acid sequence alignment of their flagellin proteins reveals that *A. tumefaciens* has a higher degree of variability in the less conserved central region. Although the amino and carboxy termini are well conserved, the internal variability suggests the possibility that flagellar filaments of *A. tumefaciens* might also have significant structural diversity (12).

In this study, we created non-polar deletions of the four flagellin genes *flaA, flaB, flaC* and *flaD* in single and all possible multiple combinations in order to genetically decipher their functions and determine the role(s) that each might play in establishing flagellar filaments in *A. tumefaciens*. We have found that the flagellins are not redundant in their functions and differ with respect to their abundance in flagella. By specific flagellin labeling, FlaA and FlaB can be visualized in otherwise normal flagellar filaments, but adopt different distributions. FlaC and FlaD are present in demonstrably lower levels. Genetic suppressor analysis has revealed that existing flagellins and other regulatory genes that affect flagellar expression can compensate for the loss of the major flagellin FlaA in flagellin mutants, and that a single key residue in FlaA is a major determinant of the helical flagellar conformation.

## Results

### FlaA is required for normal flagella synthesis and motility

Mutational analysis of the *A. tumefaciens fla* genes has been reported, identifying *flaA* gene as the only flagellin absolutely required for synthesizing a functional filament to confer normal motility, and that *flaA* mutants formed vestigial stubs (16). However, the interpretation of this work was confounded by the types of mutations generated, such as by marker-integration, which have a propensity for problems such as polar effects on downstream genes. To circumvent these complications, we created a complete set of in-frame deletions for all flagellin genes (*flaA* Atu0545, *flaB* Atu0543, *flaC* Atu0542, and *flaD* Atu0567); in all combinations (Fig. S2A and B). The Δ*flaA* mutant produces straight, somewhat shortened filaments as evaluated by transmission electron microscopy (TEM, Fig. 1A), rather than producing stubs as reported in earlier studies (15). The Δ*flaA* mutant is also quite deficient in swimming through motility agar, although less so than an aflagellate Δ*flgE* hook mutant (Fig. 1B). Other single flagellin mutants - Δ*flaB*, Δ*flaC* and Δ*flaD* make normal flagella and confer wild type swim ring diameters (Fig. 1A and 1B).

**Figure 1:**
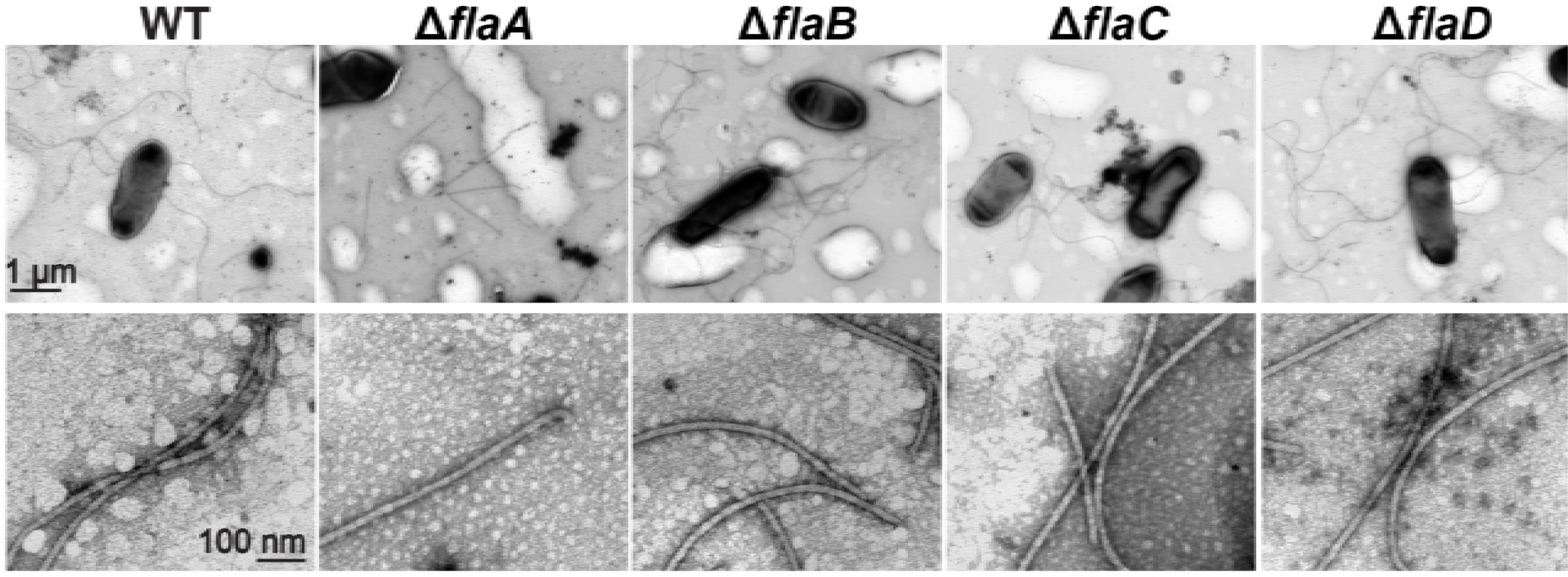
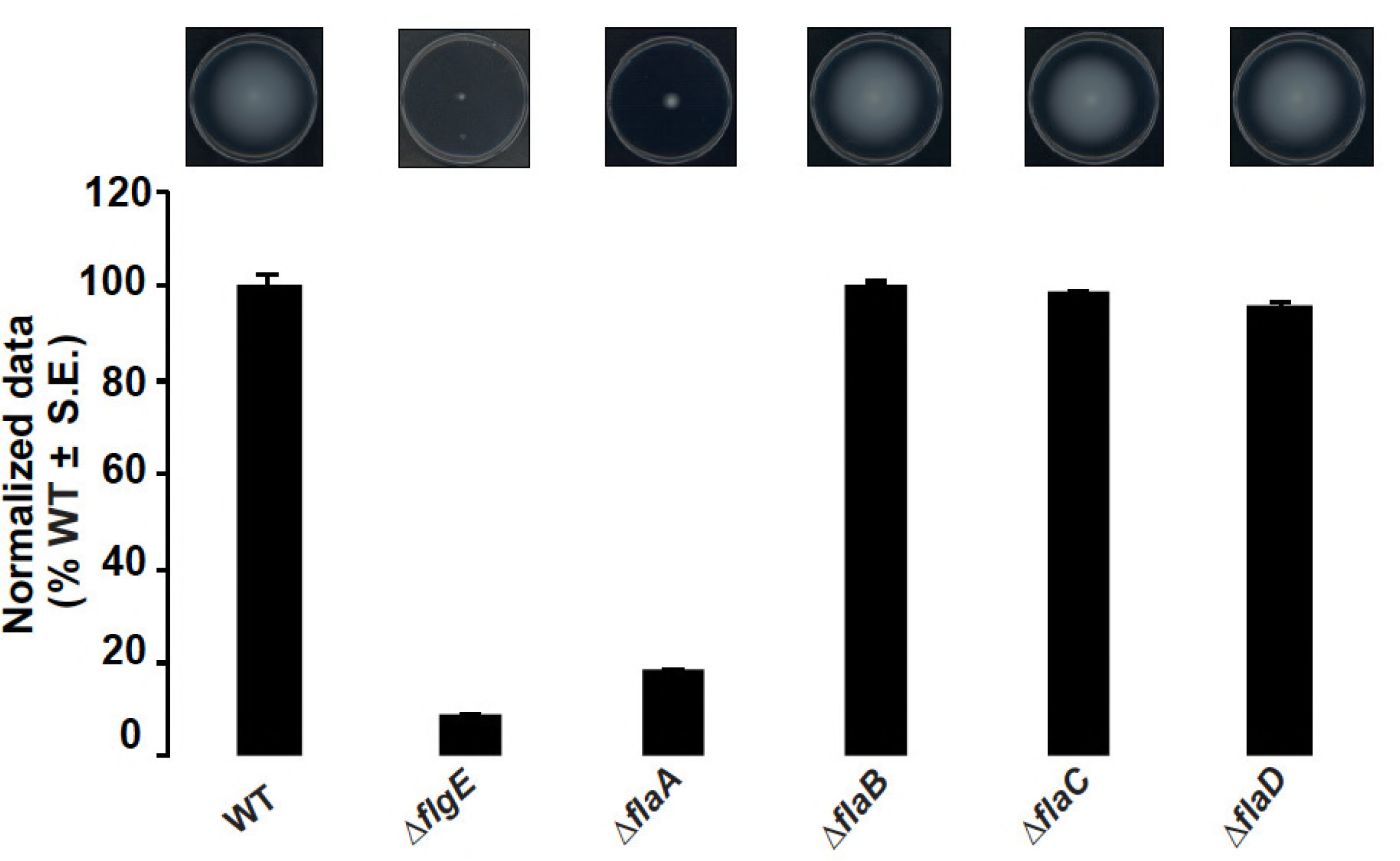
Filament morphology and swim phenotype of single deletion flagellin mutants. (A) Δ*flaA* mutants exhibit straight filaments and all other mutants: Δ*flaB*, Δ*flaC* and Δ*flaD* exhibit curved flagella similar to wild type (some sheared flagella are also shown). The top panel shows multiple filaments per mutant and the bottom panel shows zoomed in views. Images were captured at a magnification of 7500X (top panel) and 75,000X (bottom panel). Scale bar of 1 μm applies to all in top panel and scale bar of 100 nm applies to all in bottom panel. (B) Swimming motility phenotype of single deletion flagellin mutants. The top panel shows the swim plates indicating the expansion of the swim ring and the bottom panel shows swim ring diameter data (7-day incubation) for Δ*flaA,* Δ*flaB*, Δ*flaC* and Δ*flaD* flagellin mutants, all normalized to the wild type. Values are the averages for three swim plates per strain. Error bars show SEM.

All double and triple flagellin mutants with *flaA* have similar filament morphology and are equally compromised for motility (Figs. 2A, 2B). The presence of *flaB* in Δ*flaA*, Δ*flaAC*, Δ*flaAD* and Δ*flaACD* flagellin mutants correlates with longer flagella filaments than those which lack *flaB* (Δ*flaAB*, Δ*flaABC* and Δ*flaABD*; Fig. 2A), suggesting that FlaB may have a specific role distinct from FlaC and FlaD. All mutants lacking *flaA* were complemented with plasmid-borne expression of *flaA* (expressed from its native promoter in combination with the *E. coli lac* promoter, *P*_*lac*_*-P*_*flaA*_), except for the quadruple Δ*flaABCD* mutant (Fig. 2B). Additionally, the Δ*flaACD* mutant harboring the *flaA* expression plasmid was roughly 50% wild type motility, suggesting that together, the chromosomal *flaB* and the plasmid borne *flaA*, do not impart full motility.

**Figure 2:**
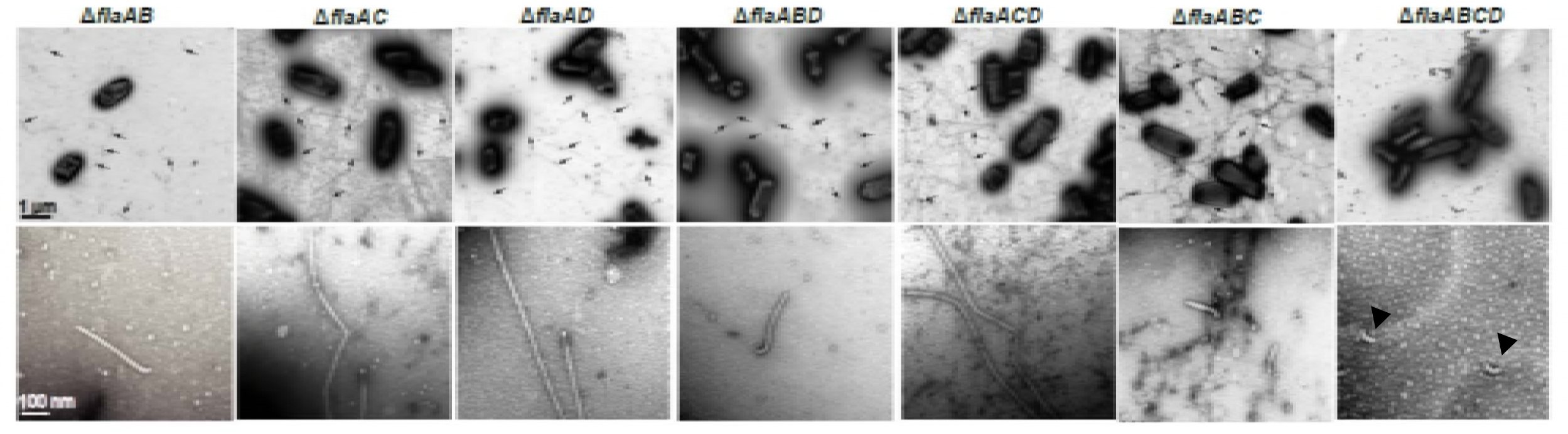
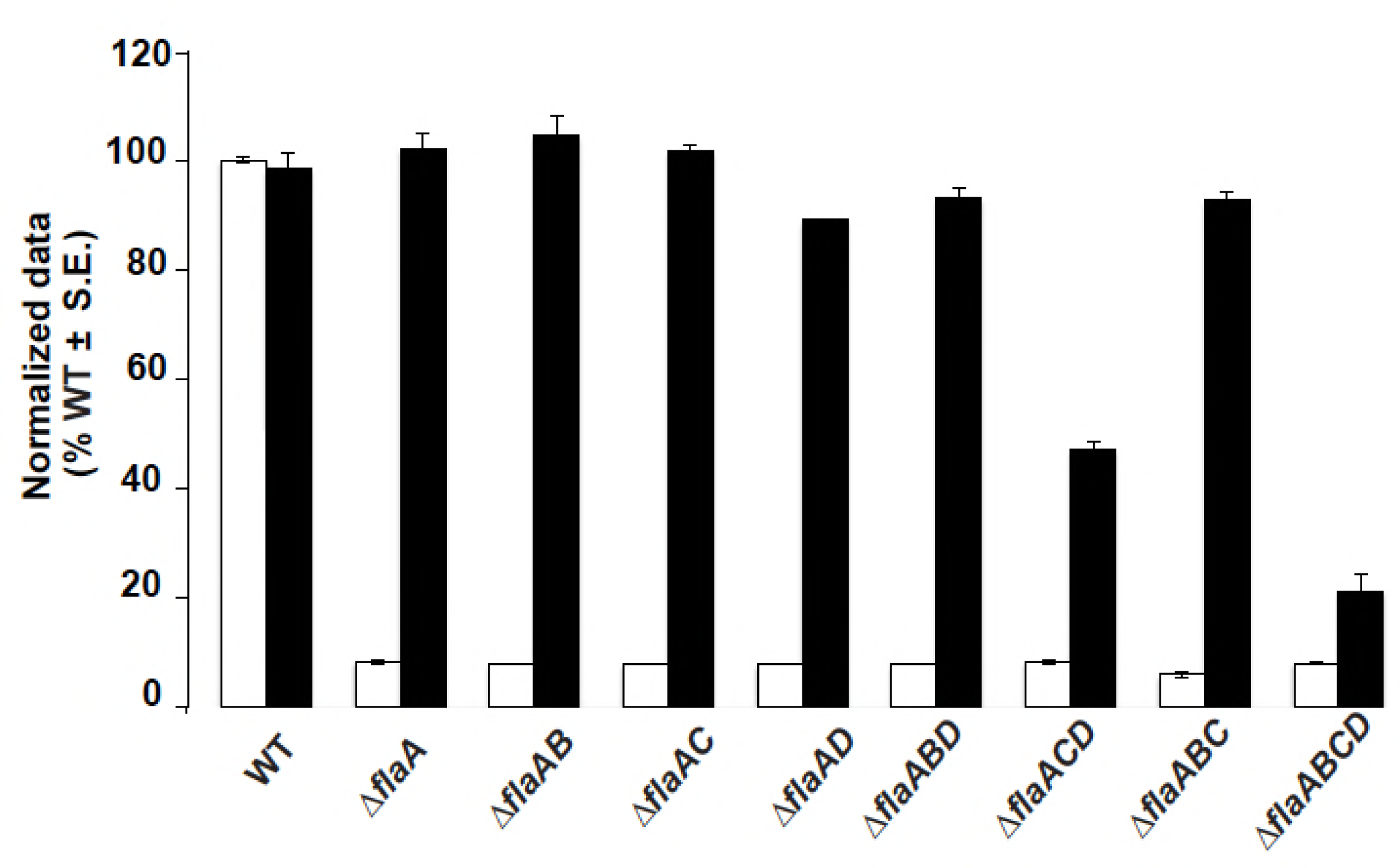
Filament morphology and swim phenotype of multiple deletion *flaA* flagellin mutants. (A) All Δ*flaA* mutants exhibit straight filaments and Δ*flaABCD* mutant is aflagellate. The top panel shows multiple filaments per mutant (indicated by black arrows) and the bottom panel shows zoomed in images of 1 or 2 filaments. Images were captured at a magnification of 6000X (top panel) and 60,000X (bottom panel). Δ*flaABCD* mutants are aflagellate and show hooks. The scale bar of 1μm applies to all images in top panel and the scale bar of 100 nm applies to all images in the bottom panel. (B) Swimming motility phenotype of all Δ*flaA* mutants harboring plasmid borne *flaA*. Swim ring diameter data (7-day incubation) for complementation of all Δ*flaA* mutants with plasmid borne expression of *flaA* expressed (*P*_*lac*_-*P*_*flaA*_-*flaA*). Data is normalized to the wild type. White bars are the wild type or original mutants and the black bars are wild type or the mutants carrying *P*_*lac*_-*P*_*flaA*_-*flaA.* Values are the averages for three swim plates per strain. Error bars show SEM.

### FlaA is required but not sufficient for proficient motility

FlaA appears to be absolutely required for biogenesis of normal flagella and motility, however it was not clear if FlaA alone is sufficient for motility. The observation that the quadruple Δ*flaABCD* mutant with plasmid borne *flaA* remained quite compromised in swimming indicated that on its own *flaA* might not be sufficient for motility (Fig. 2B). In order to investigate this directly, an in-frame deletion of the three secondary flagellins *flaB, flaC* and *flaD* was generated to obtain a Δ*flaBCD* triple mutant. The mutant exhibited very few flagellar filaments and was severely compromised in motility (Figs. 3A-C). Although the few flagella produced by the Δ*flaBCD* mutant appeared to be curved, they were still insufficient to confer normal motility. Further evidence for the requirement of these flagellins, is that the plasmid-borne *flaA* (*P*_lac_-*P*_flaA_-*flaA*) has little effect on this mutant (Fig. 3C). Dark field microscopy of cells in liquid growth media revealed some weakly swimming cells in contrast to the completely non-motile phenotype of the Δ*flaBCD* mutant alone (data not shown). This is also evident with microscopic observation of weak swimming behavior of a quadruple Δ*flaABCD* mutant with the plasmid-borne *flaA*. This suggests that elevated *flaA* expression can confer residual basal motility and its presence can create curved filaments, but that it alone does not confer normal motility.

**Figure 3:**
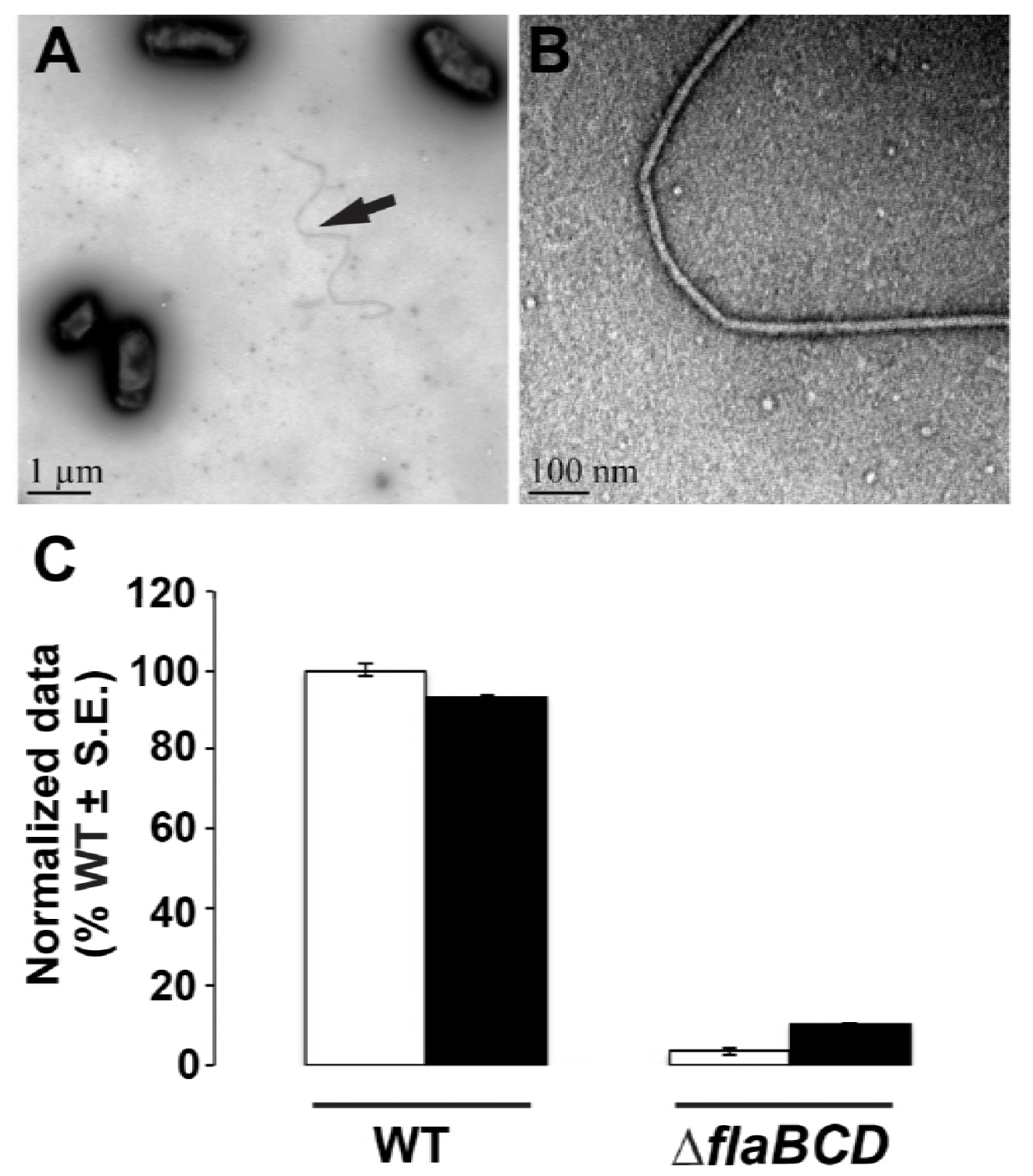
Filament morphology and swim phenotype of Δ*flaBCD* and motile flagellin mutants. (A-B) Curved filament morphology of the Δ*flaBCD* mutant. TEM images are captured at 6000X (A) and 60,000X (B). (C) Day 7 swim ring diameter data for Δ*flaBCD* mutant alone and with plasmid borne overexpression of *flaA* from *P*_*lac*_ and native promoter of *flaA* (*P*_*lac*_-*P*_*flaA*_-*flaA*). Data is normalized to wild type. White bars are wild type or the mutant and the black bars are the strains with plasmid constructs. Values are the averages for three swim plates per strain. Error bars show SEM.

### FlaA and at least one other secondary flagellin are required for curved flagellar filaments and can confer wild type motility

In order to further investigate the requirement of the *flaBCD* flagellins, we evaluated all double flagellin mutants that still retained a chromosomal copy of *flaA* along with any one of the secondary flagellins. Hence, we examined the following mutants; Δ*flaBC* (retains *flaA* and *flaD*), Δ*flaBD* (retains *flaA* and *flaC*) and Δ*flaCD* (retains *flaA* and *flaB*). All these mutants exhibited normal flagella (Fig. 4A) and near normal motility on swim agar (Fig. 4B). However, the Δ*flaCD* mutant appeared to show a significant reduction in motility in contrast to others, consistent with the observation of the Δ*flaACD* with plasmid-borne *flaA* (Fig.2B). Additionally, the Δ*flaBCD* mutant could be complemented with plasmid-borne expression of *flaB, flaC* and *flaD* indicating again that FlaA and any one secondary flagellin can confer significant levels of motility (Fig. 4B). These findings, along with the observation that the quadruple flagellin mutant Δ*flaABCD* could not be complemented with plasmid-borne *flaA* (Fig. 2B), emphasize the requirement for *flaA* and at least one of the three secondary flagellins (*flaB, flaC* or *flaD*).

**Figure 4:**
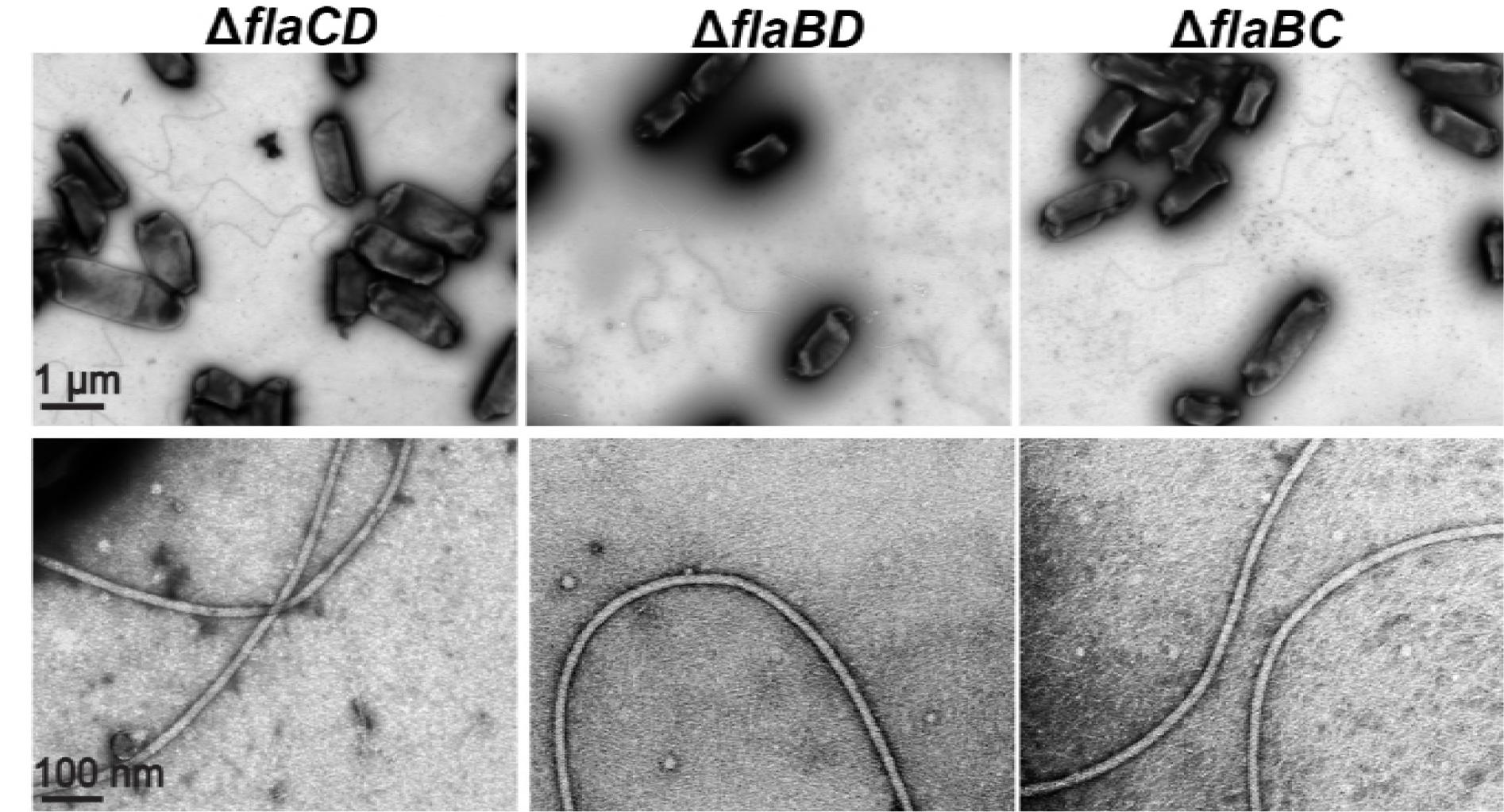
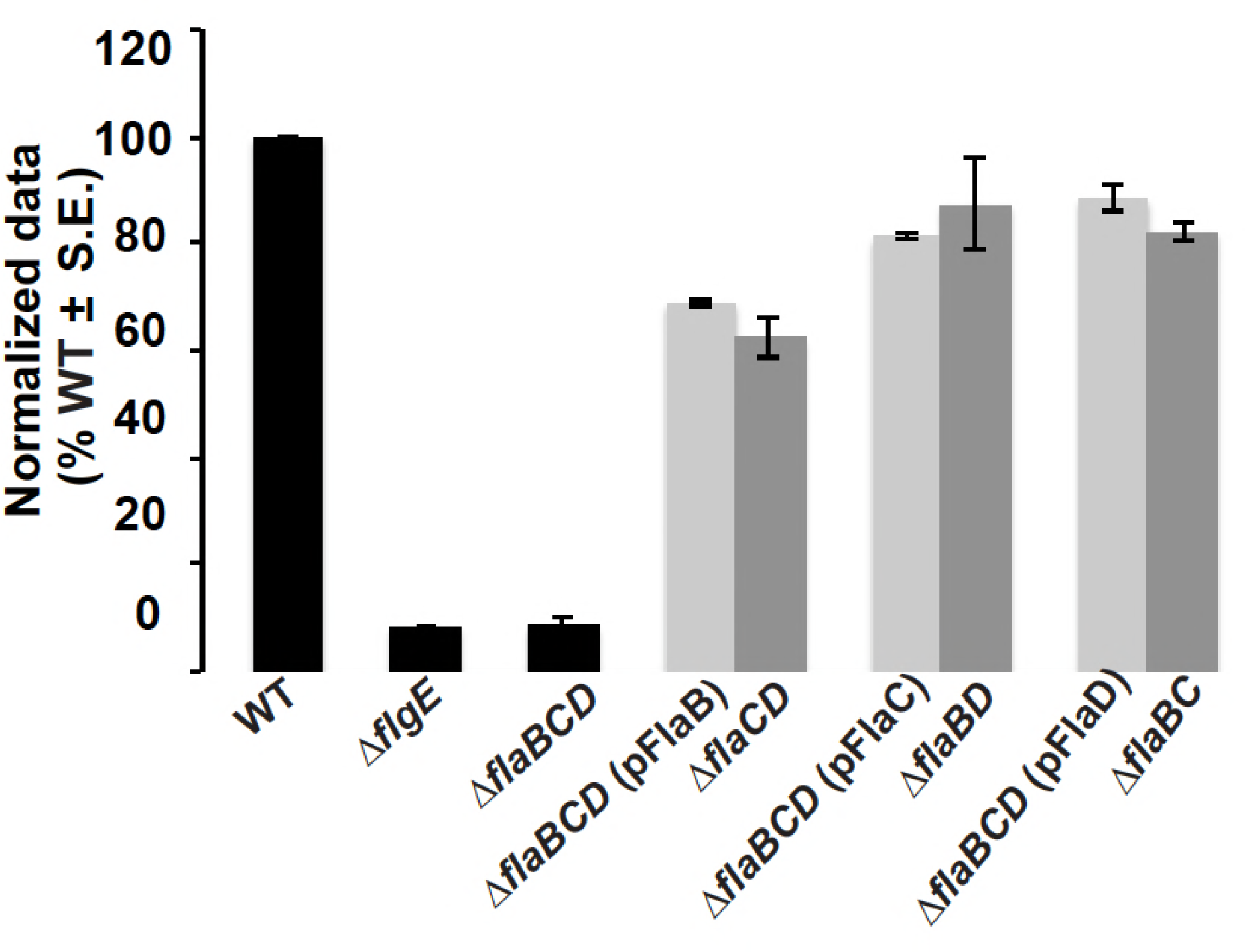
FlaA paired with any secondary flagellin restores curved flagella and motility. (A) TEM micrographs of Δ*flaCD,* Δ*flaBD and* Δ*flaBC* mutants. The top panel shows multiple filaments per mutant and the bottom panel shows zoomed in views of 1 or 2 filaments. Images were captured at a magnification of 6000X (top panel) and 60,000X (bottom panel). The scale bar of 1μm applies to all images in top panel and the scale bar of 100 nm applies to all images in the bottom panel. (B) Swim ring diameter data after 7 days for the Δ*flaBCD* mutant with plasmid borne expression of either *flaA* or *flaB* or *flaC* expressed from *P*_*lac*_ (light grey bars), induced with 400 μm IPTG. Data is normalized to the wild type. The double mutants are indicated for comparison (dark grey bars). Wild type and other controls: Δ*flgE and* Δ*flaBCD* are shown as black bars. Values are the averages for three swim plates per strain. Error bars show SEM.

### FlaA and FlaB are the most abundant flagellins in wild type filaments, but FlaC and FlaD are present at lower levels

Flagellar filament proteins of wild type and mutants had been observed by SDS-PAGE, however the proportions of flagellins in wild type filament preparations has not been reported (20). We quantified the relative amounts of the flagellins using liquid chromatography –; mass spectroscopy (LC-MS). We first prepared wild type filaments from three different replicates as described in Materials and Methods. TEM analysis confirmed the presence of intact flagellar filaments in this preparation. The preparation was incubated with trypsin, and analyzed by LC-MS. We identified signature peptides for all flagellins, FlaA, FlaB, FlaC and FlaD to be present in the wild type filament preparation. The signal intensity observed in MS is not a direct indicator of molar amount, largely due to the fact that different peptides ionize with different efficiencies. Nevertheless, unique peptides derived from identical positions in each protein will possess similar, albeit not identical, amino acid residues, and hence will ionize to similar extents. Comparing these sets of diagnostic peptides, we determined the approximate molar ratio of each flagellin (normalized to FlaB) to be FlaA (1.3), FlaB (1), FlaC (0.2) and FlaD (0.2). Examples of MS survey scans of the peptides IGLQEDFASK and IGLQEDFVSK from FlaB and FlaC, respectively are shown in Figs. 5A-B, and a plot of the signal intensities of these peptides as a function of LC elution time demonstrates the signal from the FlaB-derived peptide is much more abundant (Fig. 5C). Similarly, the MS survey scans of LVTATEEGVDR and LVAAYGVGADR, from FlaB and FlaD, respectively are shown in Figs. 5D-E, and the plot of peptide intensity as a function of time demonstrates that the FlaB peptide is the more abundant of the two (Fig. 5F). The full dataset of extracted peptide signals are reported in Table S1. Parallel analysis of the Δ*flaABC* mutant confirmed that these peptides were specific to the flagellins (data not shown).

**Fig. 5:**
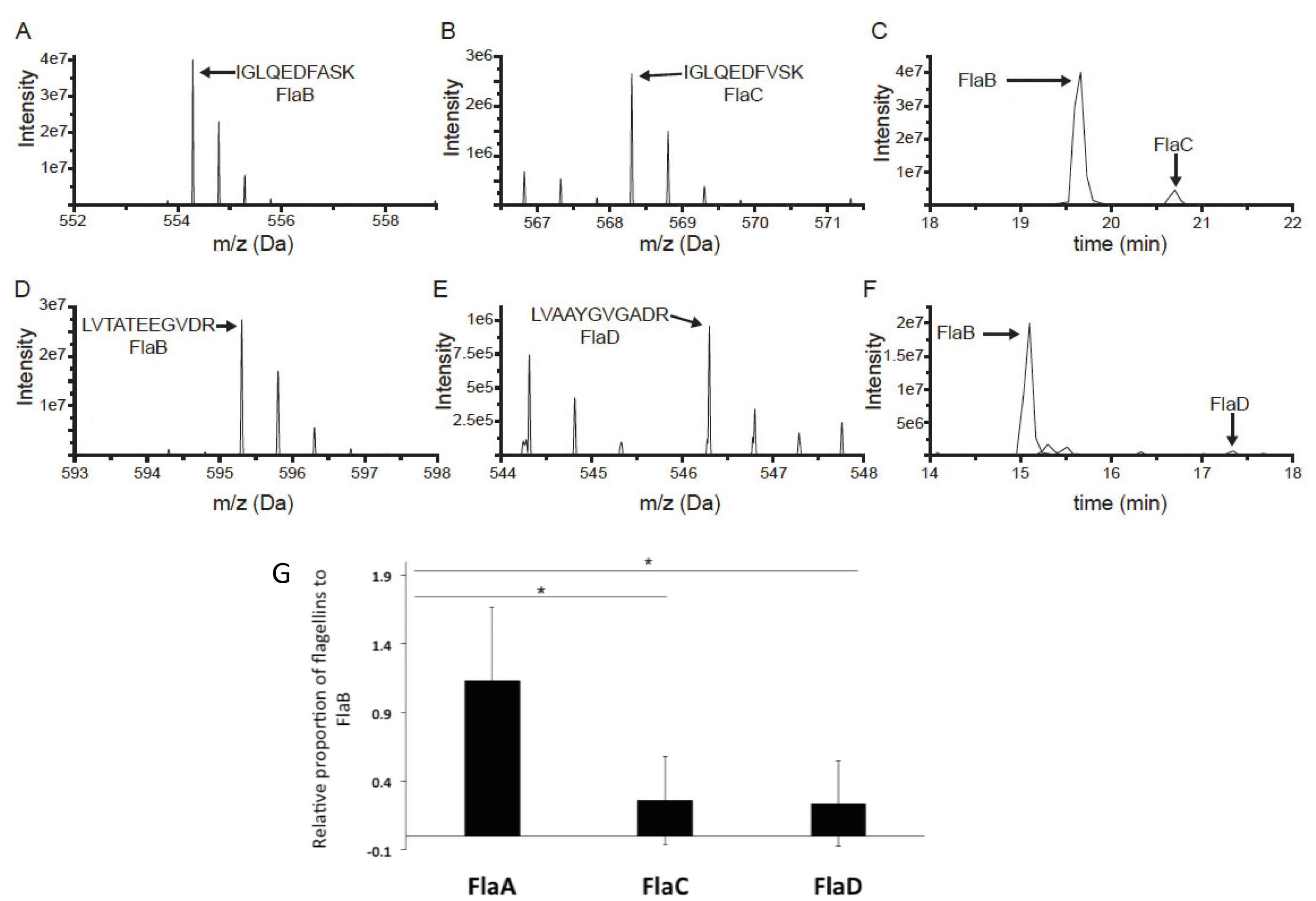
MS analysis data to find the relative proportions of flagellin proteins in wild type filament. Panels A-F are representative examples of the MS-based relative quantification of flagellin isoforms. The experiment was conducted three times, and in the case of all peptides, the identity of the MS peak was confirmed at least once by MS/MS. Figs. 5A-B shows MS survey scans of two homologous doubly-charged peptides from FlaB (IGLQEDFASK at 554.29 Da) and FlaC (IGLQEDFVSK at 568.30 Da) at the peak of their respective elution times. Fig. 5C shows the signal observed for those peptides as a function of LC elution time with the peak for the FlaB peptide approximately 10 times more intense than that of the FlaC peptide. Figs. 5D-E shows MS survey scans of two homologous doubly-charged peptides from FlaB (LVTATEEGVDR at 595.31 Da) and FlaC (LVAAYGVGADR at 546.30 Da) at the peak of their respective elution times. Fig. 5F shows the signal observed for those peptides as a function of LC elution time with the peak for the FlaB peptide approximately 25 times more intense than that of the FlaD peptide. (G) Liquid chromatography-Mass Spectrometric (LC-MS/MS) analysis reveals the relative proportions of flagellin proteins in a wild type filament preparation. Bar graph reporting the relative abundances of FlaA, FlaC and FlaD with respect to FlaB. FlaB was chosen as a reference since FlaB protein had the most peptides that were similar to peptides in the other three isoforms. The horizontal lines with an asterisk above the bars indicate that FlaA: FlaB is relatively much higher than FlaC: FlaB and FlaD: FlaB, via one-way ANOVA statistical analyses.

Comparing the ratios of different flagellin peptides and applying one-way ANOVA analysis, we found that quantities of FlaA and FlaB in a wild type filament are not significantly different. After removal of peptide ratio outliers, when normalized to FlaB levels, FlaA was significantly higher than either FlaC or FlaD. In contrast, FlaC levels were not significantly different from FlaD. This indicates that FlaA and FlaB are in higher proportions than FlaC and FlaD in a wild type *A. tumefaciens* filament preparation, and are in roughly equivalent amounts (Fig. 5G). However, observation of filament morphologies and swim assays of flagellin mutants revealed that FlaA is the functionally predominant flagellin, and that along with at least one of the other flagellins can confer proficient motility.

### Cysteine labeling of FlaA and FlaB enables visualization of flagellar filaments

In order to visualize flagellar filaments, we site specifically mutated threonine residues in the flagellin coding sequences to cysteines, allowing them to be labeled with a cysteine-reactive Alexafluor 488 maleimide dye (21). Individual flagellin genes were mutated as follows: FlaA, T213C (FlaA_Cys_); FlaB, T190C (FlaB_Cys_); FlaC, T216C (FlaC_Cys_); FlaD T276C (FlaD_Cys_). All these sites were carefully chosen so that they were in a predicted exposed region of the flagellin structure, as determined by modeling on *Salmonella* flagellin FliC with the Phyre2 program (Fig. S3A). In order to confirm these mutant alleles are functional, we tested complementation of the Δ*flaA* mutant with plasmid-borne expression of *flaA*_Cys_ and the Δ*flaBCD* mutant with plasmid-borne expression of either *flaB*_*Cys*_, *flaC*_*Cys*_ or *flaD*_*Cys*,_ all expressed from the *P*_*lac*_ promoter. All constructs complemented the corresponding mutants to the same level as their wild type counterparts indicating these constructs are fully functional (Fig. S3B). The mutant Fla_Cys_ alleles were individually exchanged for the wild type copy of the respective flagellin (Fla) in the chromosome. The site-specific mutation from threonine to _cys_teine does not hamper normal motility in these allelic replacement derivatives (Fig. S3C), although the *flaB* allele does exhibit a modest decrease in motility.

Wild type cells encoding chromosomal FlaA_Cys_, treated with Alexafluor 488 maleimide dye exhibited single curved, long, helical filaments and also bundles. FlaA seems to be present uniformly along the entire filament. Flagella are often shed by the cell (Fig. 6) as has been seen earlier in *A. tumefaciens* via transmission electron microscopy (22). In contrast to the FlaA_Cys_ mutant, cells expressing FlaB_Cys_, revealed very short flagellar fluorescent filaments, many of which were not attached to cells (Fig. 6), despite full length flagella in this mutant as indicated by TEM (data not shown) and by motility assays (Fig. S3C). This suggests that FlaB might not be present in the entire filament, but rather comprise a discrete portion of it. We did not observe any fluorescent filaments with flagellated cells expressing either FlaC_Cys_ or FlaD_Cys_, probably due to their low abundance, consistent with the LC-MS analysis (Fig. 5G).

**Fig. 6:**
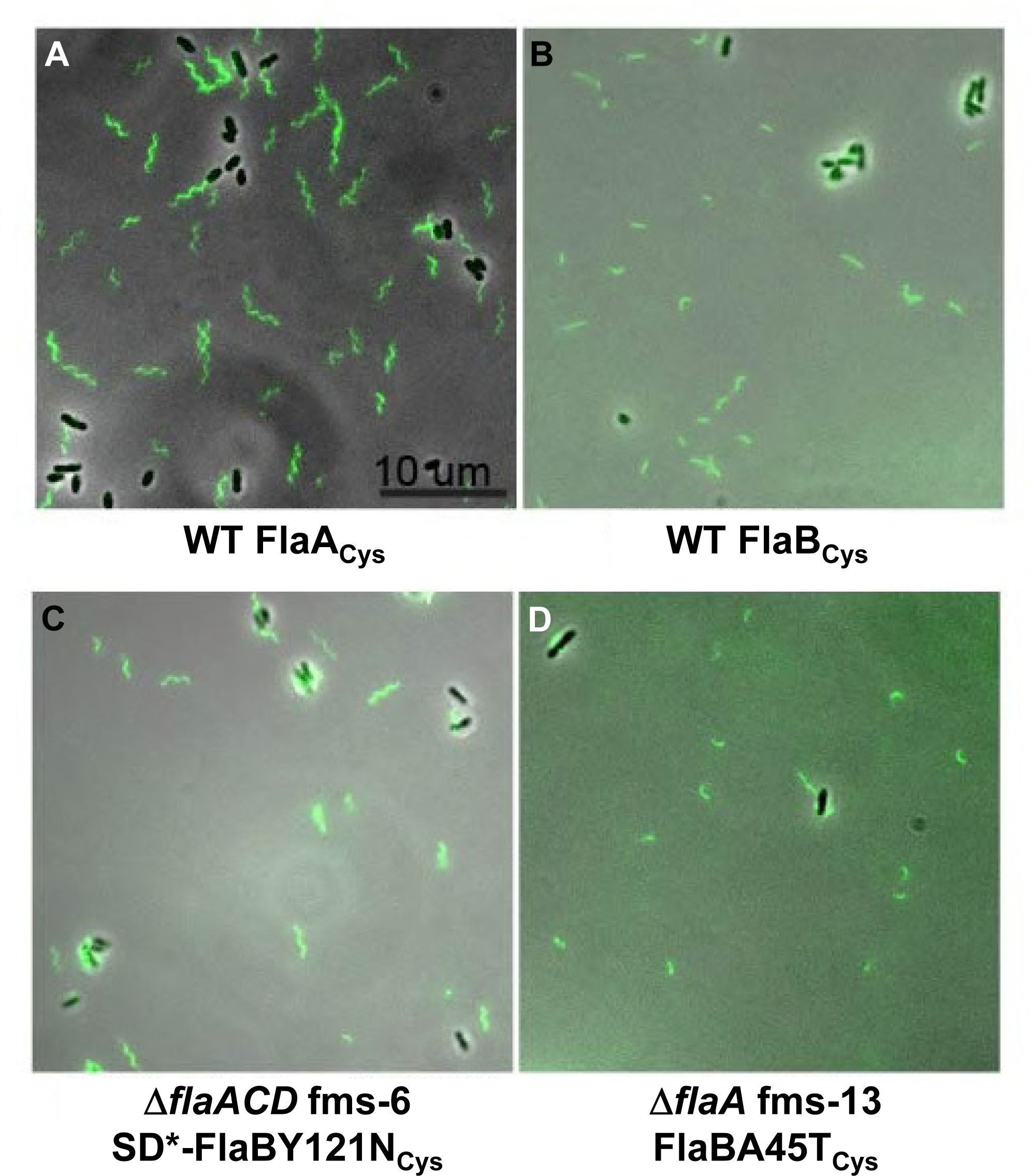
Filament morphology of wild type and flagellin mutant suppressors that have chromosomal copies of FlaA_Cys_ or FlaB_Cys_. Flagella are visualized with fluorescence microscopy of cells labeled with Alexafluor maleimide dye linkage to cysteine residues in specific flagellins. The Fla_Cys_ residues were introduced into the *flaA* (panel A) or *flaB* (panels B, C, D) by allelic exchange of the resident gene with wild type C58 (Panels A and B), Δ*flaACD* fms-6 (panel C), and Δ*flaA* fms-13. For the fms-6 and fms-13 mutants that contained both the *flaB* suppressor mutations and the cysteine allele were identified. The scale bar of 10 μm applies to all images. Flagellar filaments with cysteine labeling of FlaC and FlaD could not be visualized likely due to their low abundance. SD* refers to mutated Shine Dalgarno in the suppressor mutant. Images were collected with a Nikon Ti microscope.

### Isolation of flagellin <u>m</u>utation <u>s</u>uppressors (fms) that retain chromosomal *flaA* or *flaB* genes

Migration through motility agar is severely compromised in flagellin mutants including Δ*flaA,* Δ*flaAC,* Δ*flaAD,* and Δ*flaACD*. We observed however that extended incubation on motility agar gave rise to suppressors initially observed as flares emanating from the small rings of the parent mutants. Suppression only occurred in Δ*flaA* mutants having a chromosomal copy of *flaB*. The Δ*flaAB,* Δ*flaABC* and Δ*flaABD* mutants did not produce any flares or suppressors. The isolated Δ*flaA* suppressor mutants exhibited significantly enhanced swim rings (Fig. S4). We designated these isolates as flagellin <u>m</u>utation <u>s</u>uppressor (fms) mutants.

Several candidate fms suppressors of the Δ*flaA* mutant were selected for further analysis: Δ*flaA* fms-13, Δ*flaAC* fms-5, Δ*flaAD* fms-12, Δ*flaACD* fms-1 and Δ*flaACD* fms-6. These flagellin suppressor mutants were first evaluated by sequencing for mutations in the flagellin regulators *flaF* and *flbT*, known to control flagellin expression in *C. crescentus* and *Brucella melitensis* (23-26), however they were found to be wild type for these genes. We performed whole genome sequencing (WGS) using an Illumina platform plus further targeted DNA sequencing to identify the mutations underlying the suppression phenotypes (Table 2 and Fig. 7).

**Fig. 7:**
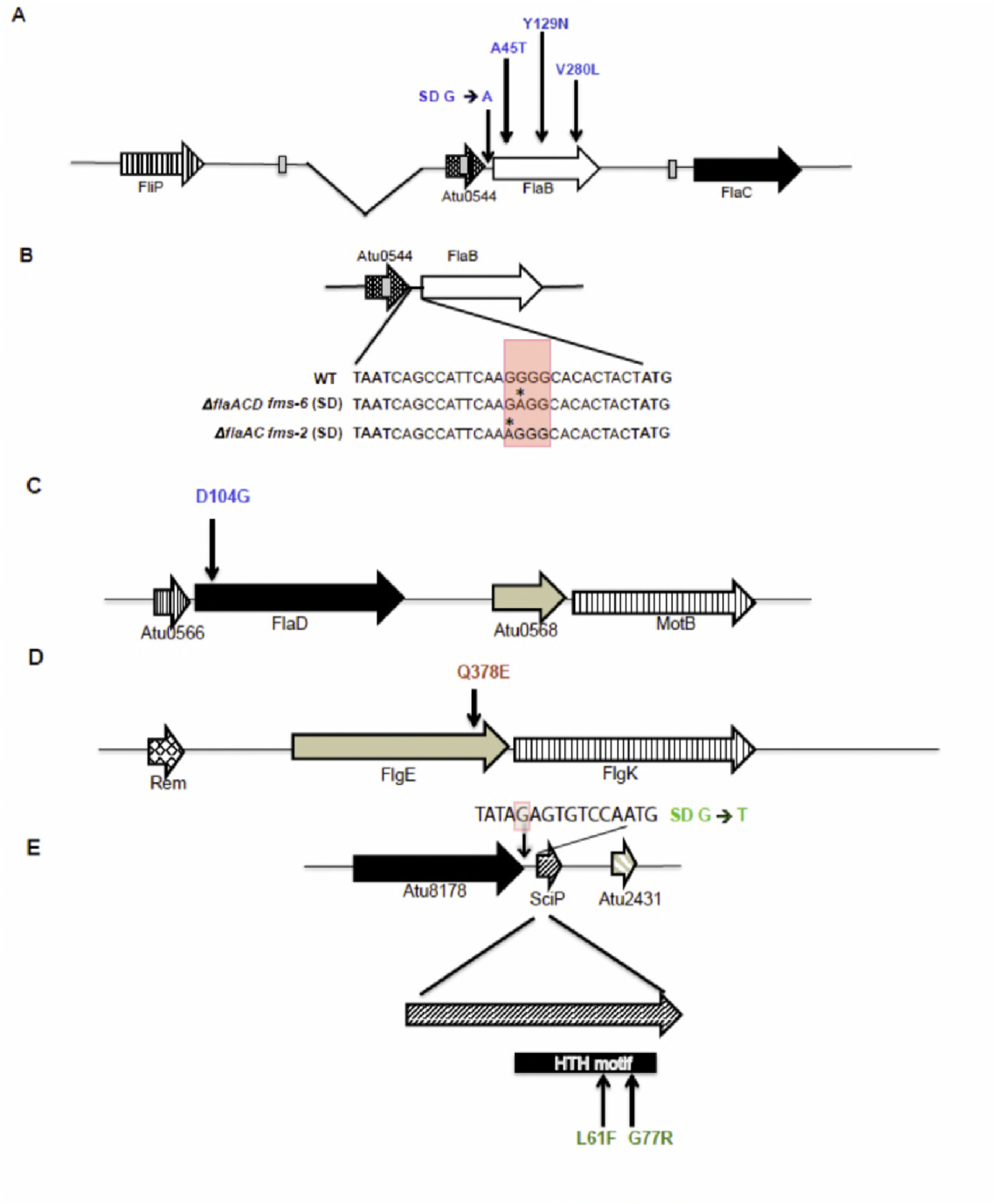
fms mutations mapped to flagellar genes and Atu2430 (SciP). The mutations are color-coded: blue for flagellin mutations, red for the hook mutation and green for SciP mutations. (A) Arrangement of *flaB, flaC*, and Atu0544 along with deletion in *flaA* indicated as the gap. Vertical arrows indicate fms mutations that are upstream of FlaB (Shine Dalgarno) and within the *flab* coding region. (B) Arrangement of Atu0544 and FlaB and the genetic sequence starting from Atu0544 stop codon (TAA) to *flaB* start codon (ATG). Details on the Shine Dalgarno mutation at genetic sequence level with red shaded box indicating predicted element and asterisk indicating the mutated residue (SD G→ A). (C-D) Arrangement of *flaD* and *flgE* respectively; vertical arrows indicate the fms mutations mapped to *flaD* and *flgE.* (E) Arrangement of SciP and with mutations mapped to Shine Dalgarno (SD* G→ T, shaded red in the sequence) and in SciP coding region. FlaB, FlaD, FlgE and SciP are 320, 430, 425, and 90 codons in length respectively.

**Table 2.** Mutations and affected region(s) in flagellin mutation suppressor (fms) derivatives.

| Mutant | Parent Mutant | WGS<sup>a</sup> | Affected Protein /Region(s) | Mutation(s) | Amino acid change(s) |
| --- | --- | --- | --- | --- | --- |
| fms-13 | $\Delta flaA$ | Yes | FlaB | G133A | A45T |
| fms-2 | $\Delta flaAC$ | No | SD<sup>b</sup>-FlaB | G>A 13 bp US of *flaB* coding sequence | No change |
| fms-5 | $\Delta flaAC$ | Yes | FlaD, FlgE, US<sup>c</sup> Atu2430 (SciP)<sup>d</sup> | A311G (*flaD*)<br>C1132G (*flgE*)<br>G>T 9 bp US (Atu2430) | D104G (FlaD)<br>Q378E (FlgE) |
| fms-12 | $\Delta flaAD$ | Yes | FlaB, Atu2430 (SciP)<sup>d</sup> | G838C (*flaB*)<br>C181T (Atu2430) | V280L (FlaB)<br>L61F (Atu2430) |
| fms-1 | $\Delta flaACD$ | Yes | FlaB, Atu2430 (SciP)<sup>d</sup> | T385A (*flaB*)<br>G229C (Atu2430) | Y129N (FlaB)<br>L61F (Atu2430) |
| fms-6 | $\Delta flaACD$ | Yes | FlaB and SD<sup>b</sup>-FlaB | T385A (*flaB*)<br>G>A 12 bp US of *flaB* start codon | Y129N |

<sup>a</sup> WGS – whole genome sequence determined
<sup>b</sup> SD – Shine Dalgarno
<sup>c</sup> US – upstream
<sup>d</sup> Atu2430 is a homologue of SciP

The most common mutations identified among the suppressors were either in the *flaB* coding sequence or its upstream sequences, often in combination with additional mutations affecting motility. We found that Δ*flaA* fms-13 had a single base substitution (G133A) in the *flaB* flagellin gene (Atu0543) producing a mutant FlaBA45T protein. We characterized two mutants isolated from the Δ*flaAC* parent. The Δ*flaAC* fms-2 suppressor had a distinct G→A substitution located 13 bp upstream of the *flaB* coding sequence (Table 2), and thus had incurred a base substitution in its predicted Shine-Dalgarno sequence (Fig. 7A). However, this mutant was not analyzed by WGS and hence we are uncertain if there are other contributing mutations.

We performed WGS on the Δ*flaAC* fms-5 suppressor, and in contrast to the two mutants described above, found it did not have mutations in or around *flaB*. Rather fms-5 had three mutations (Table 2) including (i) a single base substitution (A311G) in the *flaD* flagellin gene (Fig. 7C), producing FlaDD104G mutant protein, (ii) and a second mutation (C1132G) in the hook gene *flgE* (Fig 7D; Atu0574, 1278 bp), producing a mutated FlgEQ378E protein, and (iii) a third mutation creating a single base substitution (G→T) located 9 bp upstream in the Shine Dalgarno region of the predicted transcription factor *sciP* (Fig. 7E; Atu2430, 276 bp). Originally identified in *C. crescentus* (27), SciP (small CtrA inhibitor protein) is an essential helix-turn-helix (HTH) transcription factor, and has been shown to regulate several flagellar and chemotaxis genes (28).

The Δ*flaAD* fms-12 suppressor was found to have incurred two mutations, a single base substitution (G838C) in the *flaB* flagellin gene producing FlaBV280L (Fig. 7A), and a second single base substitution also in Atu2430 (C181T), encoding the *sciP* regulator generating a predicted SciPL61F mutant protein. The Δ*flaACD* triple mutant was particularly interesting as it only has the single flagellin gene *flaB*, and thus we analyzed two independent suppressor mutants from this parent. The Δ*flaACD* fms-1 suppressor had a single base substitution (T385A) in the *flaB* flagellin gene producing FlaBY129N (Table 2), along with another single base substitution (G229C) also in the *sciP* gene (Atu2430), yielding SciPG77R (Fig. 7E). The Δ*flaACD* fms-6 suppressor had the identical Y129N FlaB flagellin mutation as with fms-1, along with another single base substitution (G→A) located 12 bp upstream of the *flaB* coding sequence (Table 2) in its predicted Shine-Dalgarno sequence, which we designate as SD*-FlaB (Fig. 7B).

All of the mutated flagellin residues are conserved between flagellins (FlaBA45, FlaBV280 and FlaDD104) with the exception of FlaBY129 (Fig. S1). Both SciP mutations L61 and G77 mapped to its HTH motif and are conserved between SciP from *A. tumefaciens* and *C. crescentus* (Fig. S5).

### Recreation of fms suppressor mutations to test sufficiency

Allelic replacement was performed for the wild type genes in the parent strains with the mutated copy of the implicated fms allele. The fms mutations were recreated either as a one-step allelic replacement as for Δ*flaA* fms-13 where only *flaB* was mutated or in a multi-step process as for Δ*flaAD* fms-12 where *flaB* and *sciP* were mutated (see Materials and Methods). Recreated Δ*flaA* fms-13 and Δ*flaA*CD fms-6 mutants restored effective motility equivalent to the respective fms mutant, when either mutated *flaB* (for fms-13) or mutated *flaB* along with its Shine-Dalgarno mutation (for fms-6) were replaced in their parent backgrounds (Fig. 8). The recreated Δ*flaAC* fms-5, Δ*flaAD* fms-12 and Δ*flaA*CD fms-1 however, could not completely restore the suppression swim phenotype with single allelic replacements of either mutated *flaB* (for fms-12 and for fms-1) or *flgE* (for fms-5) alone. Both of the recreated Δ*flaAD* fms-12 and Δ*flaA*CD fms-1 also needed their respective *sciP* alleles in order to completely restore the suppression phenotype. Interestingly, recreated Δ*flaAC* fms-5 with the mutated *flgE* allele alone, did not rescue the non-motile phenotype of Δ*flaAC* at all. Motility was partially restored to about the same degree with the mutated *flaD* allele alone or in combination with *flgE* mutation. It is possible that recreated Δ*flaAC* fms-5 will be completely restored if the region upstream of its *sciP* is also replaced with the mutated allele, since in other cases (for fms-1 and fms-12), mutated *sciP* appears to play an important role in the suppression phenotype (Fig. 8).

**Fig. 8:**
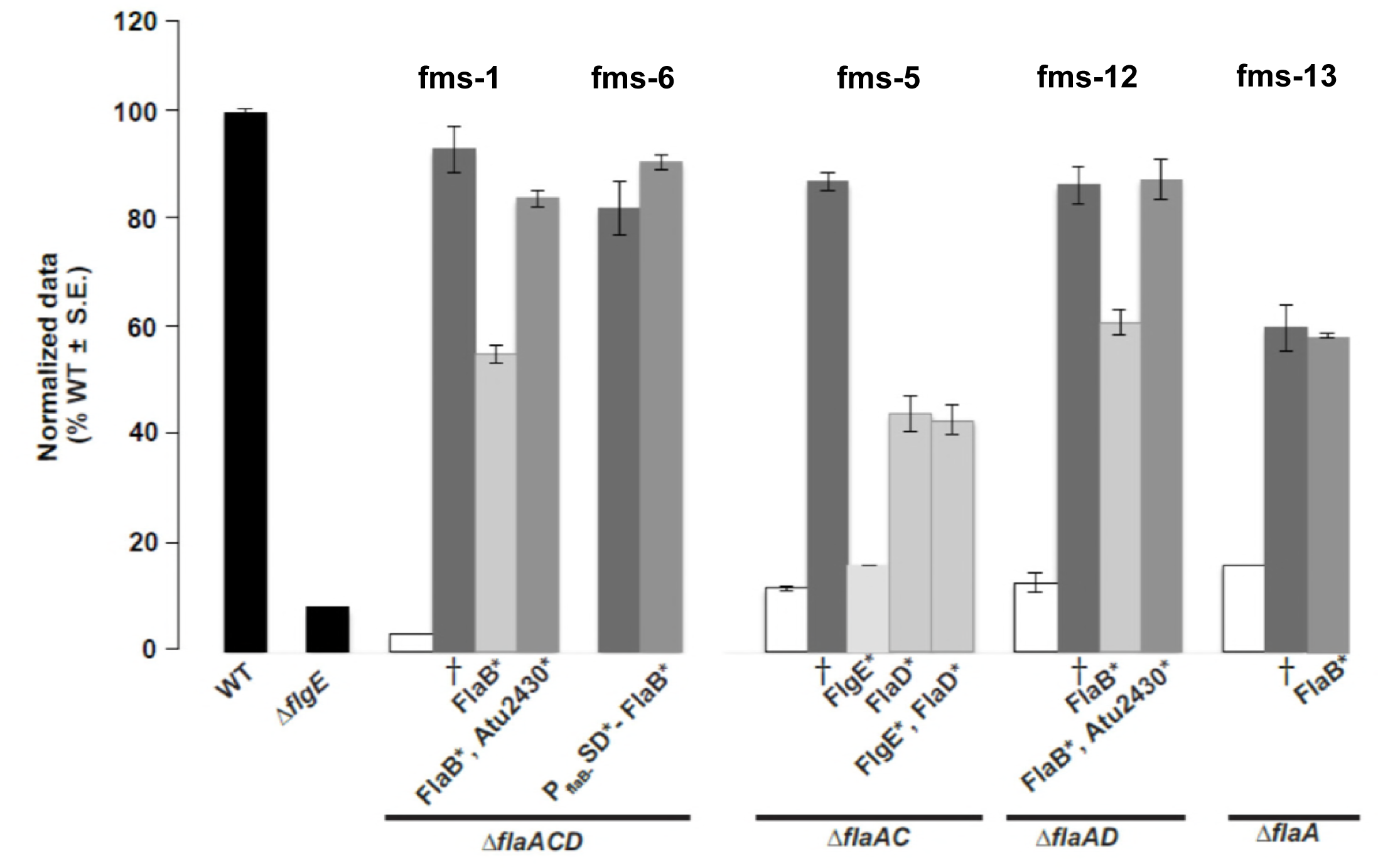
Swimming motility phenotype of fully and partially recreated fms mutants. Swim ring diameter data after 7 days showing the motility phenotype for originally isolated fms mutants and their recreated counterparts. The relevant fms mutant for each set of bars is indicated above them. Data is normalized to the wild type. Wild and Δ*flgE* are shown as black bars. White bars indicate the parent mutants. Dark grey bars are the originally isolated fms mutants (†). Light grey bars indicate the recreated suppressors that do not completely phenocopy the original suppressor. Medium grey bars indicate recreated mutants that completely reproduce the original fms phenotype. Asterisk (*) by the protein names indicate the mutated residues incorporated via allelic replacement in those strains. Values are the averages for three swim plates per strain. Error bars show SEM.

### FlaBY129N mutation can confer helicity to FlaB flagellar filaments

FlaA and FlaB were both found by LC-MS to be in similar proportions in a wild type flagellar filament preparation (Fig. 5G). Since the fms suppressor mutations commonly mapped to FlaB, we further examined how the mutated FlaB might confer suppression and enhance motility. Also, all Δ*flaA* mutants make straighter, non-helical and non-functional flagella (Figs. 1A & 2A) but the suppressors have restored motility and thus either reverse or overcome the conformational deficiency. We introduced by allelic replacement the FlaB_Cys_ mutation into two suppressor mutants that had been confirmed to require only mutated FlaB (Fig. 8) to recreate efficient motility; Δ*flaA* fms-13 that was mutated for FlaBA45T and Δ*flaACD* fms-6 that had both SD*-FlaB and FlaBY129N mutations (Fig. 7). The Δ*flaA* fms-13 - FlaB_Cys_ mutant exhibited short curved flagellar filaments very similar to WT with FlaB_Cys_, but interestingly, Δ*flaACD* fms-6 FlaB_Cys_ exhibited helical flagellar filaments very similar to those observed for WT FlaA_Cys_ (Fig. 6C and 6D). Hence both these suppressors made functional flagella but seemed to do so by different mechanisms.

It was however, unclear which mutation in Δ*flaACD* fms-6 (SD*-FlaB or FlaBY129N) contributed to the helical conformation of the FlaB-labeled flagella filament. In order to ascertain which mutation was responsible, we separated these two mutations. We first compared the expression levels of a plasmid-borne wild type *P*_*flaB*_-SD-*lacZ* translational fusion with that from a mutated *P*_*flaB*_*-*SD\**-lacZ* (with the fms-6 mutation) in Δ*flaACD* mutant via a β-galactosidase assay. The expression for the mutated Shine Dalgarno of *flaB* is higher compared to the wild type (Fig. S6). It therefore seems likely that the SD*-FlaB mutation increases FlaB production.

We had observed repeatedly that plasmid-borne *flaA* expression driven from *P*_*lac*_ in combination with the native promoter of *flaA* (*P*_*fla*__*A*_) complements all Δ*flaA* mutants entirely, in contrast to expression from *P*_*lac*_ alone in a Δ*flaA* mutant, presumably reflecting the strength of the *P*_*flaA*_ promoter (Compare Fig. 2 with S3B). We first expressed FlaB_Cys_ from the plasmid as a fusion to *P*_*lac*_*-P*_*flaA*_ and examined this construct in the Δ*flaA* and Δ*flaACD* mutants. The Δ*flaA* mutant was included to examine the effect of the plasmid-borne FlaB in a background where other flagellins were also present, and was observed to partially rescue motility, whereas its expression in the Δ*flaACD* mutant did not (Fig. 9A). Both Δ*flaA* and Δ*flaACD* with *P*_*lac*_ *-P*_*flaA*_*-*FlaB_Cys_ exhibited straight filaments (Fig. 10 E-F) that correlate with their decreased swim ring diameters. We next expressed FlaBY129N_Cys_ from *P*_*lac*_ *-P*_*flaA*_ and both Δ*flaA* and Δ*flaACD* were fully rescued for motility and exhibited helical flagella indistinguishable from the *P*_*lac*_ *-P*_*flaA*_*-*FlaA_Cys_ constructs in the same mutants (Fig. 9A, 10 A-B, G-H). This suggests that FlaBY129N is important for formation of helical flagella that impart proficient swimming in the suppressor mutants.

**Fig. 9:**
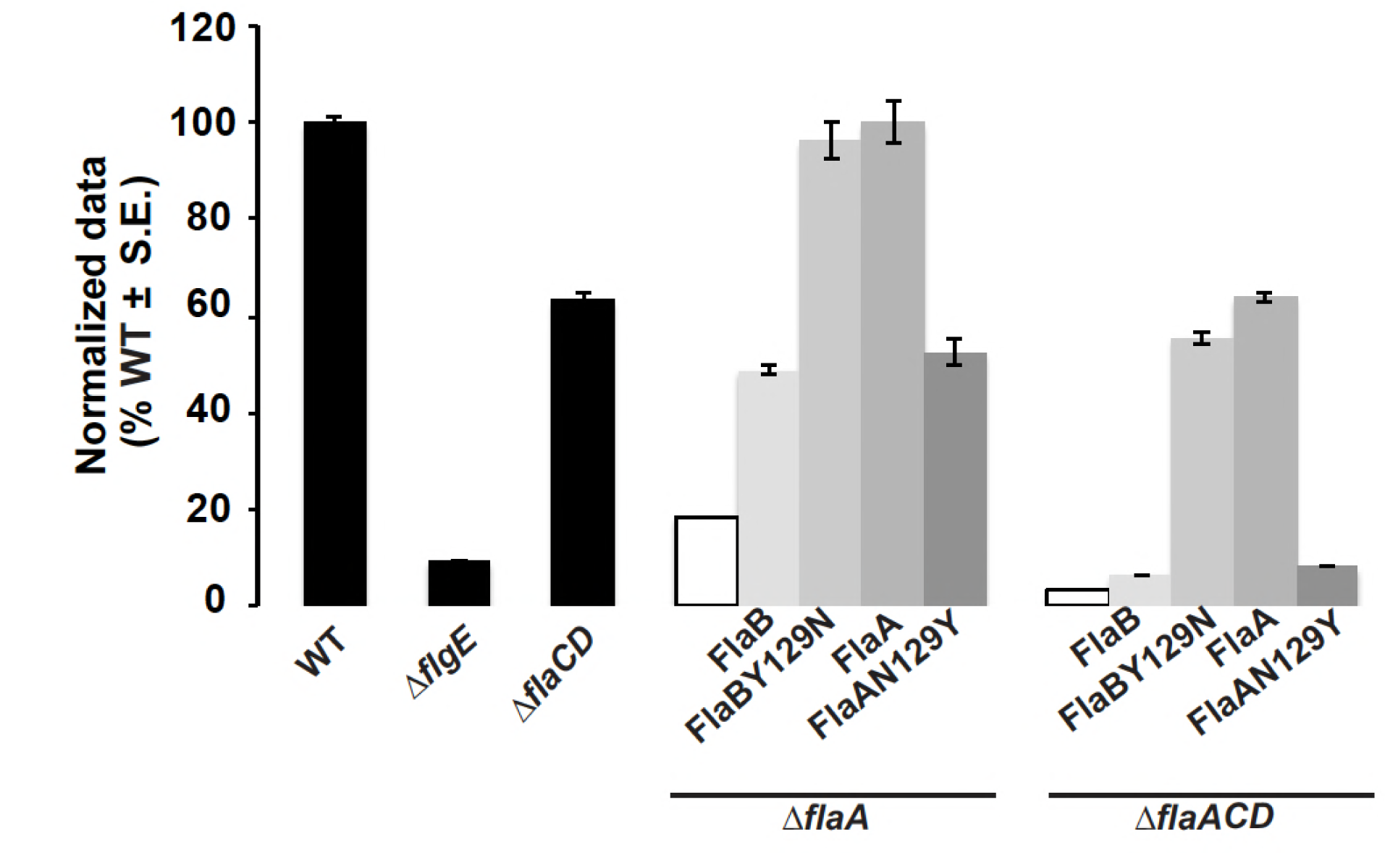
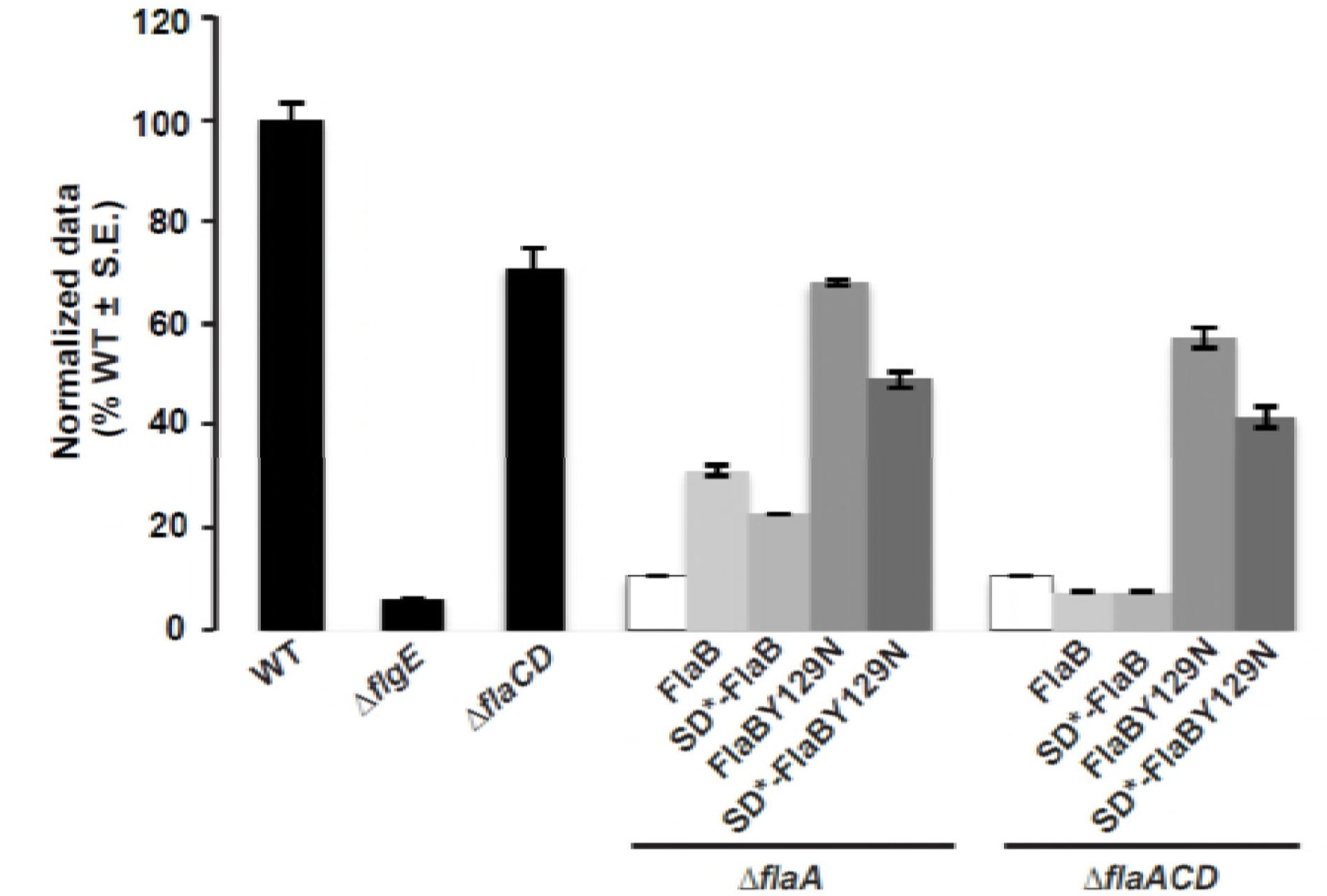
Swimming motility of Δ*flaA* and Δ*flaACD* mutants harboring plasmids carrying *flaA* and *flaB* alleles that impart altered helicity. Swim ring diameter of mutant *A. tumefaciens* derivatives. Wild type, Δ*flgE* and Δ*flaCD* are provided (black bars) for reference. The Δ*flaA and* Δ*flaACD* mutants (white bars), and derivatives harboring plasmids expressing FlaB_Cys_ (light grey) or FlaBY129N_Cys_ (light medum grey) and FlaA_Cys_ (dark medium gray) or FlaAN129Y_Cys_ (Dark grey) expressed from *P*_*lac*_*-P*_*flaA*_ (panel A) or from *P*_*lac*_-*P*_*flaB*_ as well as the SD* mutation in the flaB Shine-Dalgarno sequence (panelB). Data is normalized to the wild type. Values are the averages for three swim plates per strain. Error bars show SEM.

**Fig. 10:**
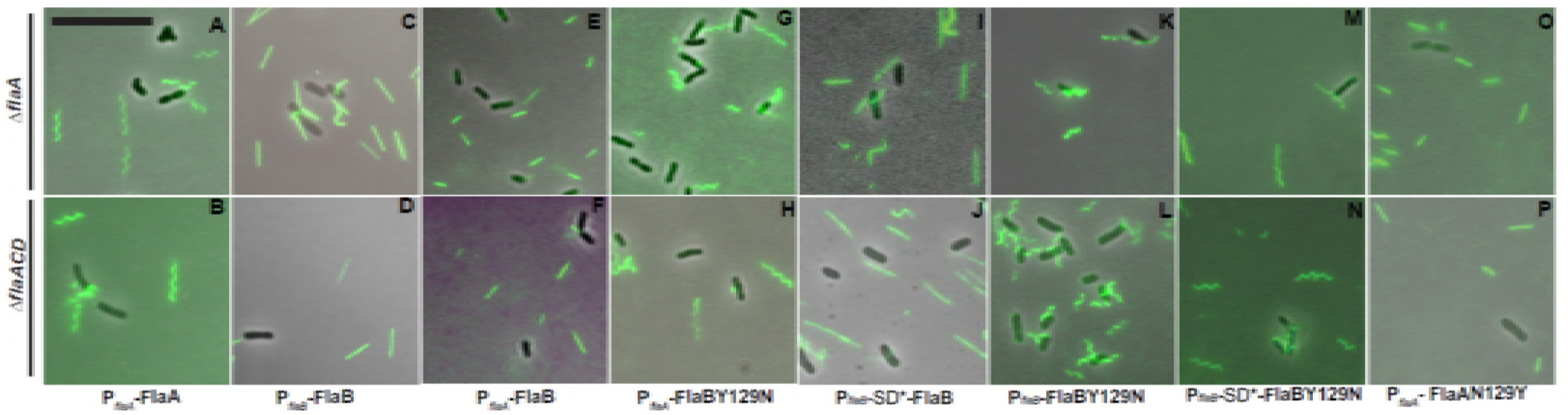
Filament morphology of plasmid-borne flagellin mutants visualized via fluorescence microscopy. Alexafluor 488 maleimide labeling of *A. tumefaciens* derivatives from Δ*flaA* (top row) and Δ*flaACD* (bottom row) – flagellin constructs are expressed from *P*_*lac*_*-P*_*flaA*_ or *P*_*lac*_*-P*_*flaB*_ (as indicated). Mutants express either FlaA_Cys_ or FlaB_Cys_ or SD*-FlaB_Cys_ and FlaAN129Y_Cys_ or FlaBY129N_Cys_ as indicated. Scale bar indicated is 10 μm and applies to all images.

In order to examine the impact of the *flaB* Shine Dalgarno mutation (from the fms-6 mutant), we expressed the wild type FlaB_Cys_ allele and the mutant derivatives, SD*-FlaB_Cys_, FlaBY129N_Cys_ and SD*-FlaBY129N_Cys_ from *P*_*lac*_ *-P*_*flaB*_ in Δ*flaA* and Δ*flaACD* mutants and observed their flagella and swimming proficiency. Both mutants expressing plasmid borne FlaB_Cys_ constructs exhibited mostly straight flagella with occasional curvature similar to WT FlaB_Cys_ (Fig. 10C and D) and only Δ*flaA* mutant showed a modest improvement in motility (Fig. 9B). Cells with SD*-FlaB_Cys_ constructs also displayed a similar phenotype as observed with FlaB_Cys_ (Fig. 10I and 10J). Interestingly, cells expressing FlaB_Cys_Y129N or SD*-FlaBY129N_Cys_ from *P*_*lac*_ *-P*_*flaB*_ exhibited helical flagellar filaments, with flagella for FlaBY129N_Cys_ (Fig. 10 K and 10L) that are qualitatively shorter that for the SD*-FlaBY129N_Cys_ (Fig. 10M and 10N). This suggests that FlaBY129N is independently contributing to helical flagella and that the Shine Dalgarno mutation is contributing by increasing flagellin levels synthesis. Both Δ*flaA* and Δ*flaACD* mutants could be partially rescued with *P*_*lac*_ *-P*_*flaB*_ driven expression of either FlaBY129N_Cys_ or SD*-FlaBY129N_Cys_ (Fig. 9B). Here, we do observe a slight decrease in swim ring diameter for both the mutants harboring SD*-FlaBY129N_Cys_ (fms-6 mutation) constructs, consistent with a slight reduction of motility with the FlaB_Cys_ allelic replacement in WT (Fig. S3C), suggesting that this allele may manifest a minor deficiency.

### Asparagine at position 129 in FlaA contributes to the formation of helical flagella

Our results suggest that the FlaBY129N mutation is a major determinant for helical flagella in the Δ*flaACD* fms-6 suppressor, and also when this allele was expressed from a plasmid in both Δ*flaA* and Δ*flaACD* mutants. We therefore examined the function of this residue in the major essential flagellin FlaA. In FlaA the residue at this position within the projected flagellin structure (Fig. S3A) is an asparagine (N129), corresponds to the tyrosine (Y129) in FlaB (Fig. S1) that is mutated to an asparagine in several suppressor mutants. In order to investigate whether N129 in FlaA is important for rendering helical flagellar filaments, we site-specifically mutated FlaAN129_Cys_ to FlaAY129_Cys_ and expressed it from a plasmid borne *P*_*lac*_*-P*_*flaA*_ in Δ*flaA* and Δ*flaACD* mutants. Interestingly, both the Δ*flaA* and the Δ*flaACD* mutantz expressing *P*_*lac*_*-P*_*flaA*_-FlaAN129Y_Cys_ exhibited straighter flagella (Figs. 10A-B, 10O-P) and significantly weaker complementation (Fig. 9A/9B) in contrast to the helical flagella and full complementation displayed by a mutant expressing FlaA_Cys_. This indicates that FlaAN129 is indeed important for formation of helical flagella and thus an important contributor to motility.

## Discussion

Bacteria synthesize flagellar filaments from either one or multiple flagellins, and it is thought that these differences might have evolved in response to the environmental niche of each particular bacterial taxon (6, 29-31). Complex flagellar filaments comprised of multiple flagellins are thought to have evolved in soil bacteria like *S. meliloti* and *Agrobacterium* sp. H13-3 to provide stable filaments that enable efficient swimming and propulsion in highly viscous environments (6, 31). For host-associated bacteria, flagella are often recognized by the host defense response as pathogen-associated molecular patterns (PAMPs) (32). In fact, members of the Rhizobiaceae, with complex flagella lack the recognized flagellin motif and are instead recognized by the host through perception of the translation factor Ef-Tu (33). It is clear that flagella are under multiple levels of strong selection in the environment for their abundance, distribution and composition.

During flagellum assembly, flagellin proteins are transported through the narrow lumen of the filament and added at the growing tip. Recently, transport and assembly has been proposed to proceed via an injection-diffusion mechanism (34). Thus, the entry of flagellins into the export channel is a major step in the process. Given the proposed injection-diffusion assembly, the mechanism by which bacteria coordinate flagella biosynthesis with multiple flagellins is even more intriguing.

In this study, we have systematically examined the function of the four flagellins that are thought to comprise the lophotrichous flagella tuft of *A. tumefaciens*. In certain bacteria such as *C. crescentus*, multiple flagellins are thought to be redundant (11), and in others they are differentially regulated or have unique functions (12, 14, 18, 19, 35, 36). In *A. tumefaciens*, FlaA is thought to be the major flagellin with accessory roles for the secondary flagellins – FlaB, FlaC and FlaD. Δ*flaA* mutants were thought to form vestigial stubs and were impaired for motility (16, 20). However, our studies with carefully constructed, in-frame mutations reveal that rather than forming stubs, Δ*flaA* mutants form filaments that lack the curvature found in wild type filaments. The Δ*flaA* mutants do not appear to be completely immobilized, as for the hookless Δ*flgE* mutant (Figs. 2A & B) and under the light microscope very slow wiggling or traversing the field of view is observed (data not shown). Our findings suggest that FlaA is the major flagellin in part because it engenders helical shape to the filament.

The role of secondary flagellins was not clear from earlier studies. However, in the related *Sinorhizobium meliloti* and *Agrobacterium* sp. H13-3 secondary flagellins are also required for proficient swimming motility: Bacteria producing FlaA together with any one secondary flagellin are able to assemble a functional filament and thus enable the cells to swim (12). We performed detailed analysis in *A. tumefaciens* to ascertain the function of individual flagellins.

We found that all triple flagellin mutants: Δ*flaBCD*, Δ*flaABC,* Δ*flaABD* and Δ*flaACD* are capable of synthesizing filaments. In other words, every flagellin (FlaA, FlaB, FlaC or FlaD) can form filaments regardless of being a major or secondary flagellin. However, the shape and lengths of the filaments varied. FlaA alone (Δ*flaBCD* mutant) forms curved filaments and without FlaA the curvature is lost, almost certainly because of loss of the molecular interactions that stabilize helical shape (Fig. 2A, 3A-B). Although all other mutants with single flagellins can synthesize a filament alone, it appears that FlaB alone (Δ*flaACD* mutant) synthesizes longer filaments than FlaC (Δ*flaABD* mutant) or FlaD (Δ*flaABC* mutant). Mutants with FlaD alone make the smallest filaments, while those with FlaB alone make notably long filaments (Fig. 2A). It is possible that all four flagellins may be represented in wild type filaments, with FlaA and FlaB as the most abundant, followed by FlaC and finally FlaD. Interestingly, this is evident from our LC/MS-MS analysis (Fig. 5G) and visualization of flagella via _cys_teine labeling of individual flagellins where only FlaA and FlaB are visible, and FlaC and FlaD are not (Fig. 6). It is not possible to comment conclusively regarding the order of flagellin assembly because we find FlaA and any one secondary flagellin (including the most divergent of the three, FlaD) are able to assemble a functional filament. It is interesting however that a Δ*flaCD* mutant is modestly compromised in swimming in comparison to other double flagellin mutants, even though they appear to have curved flagella (Figs. 4A & B). The decrease in motility might be due to an unknown accessory function that FlaC and FlaD provide, and that FlaB cannot accomplish as efficiently. We observe short curved stubs in the aflagellate Δ*flaABCD* mutants, which may be shed hooks (Fig. 2A). A small number cells were observed to be weakly swimming under the microscope (data not shown) when Δ*flaBCD* and Δ*flaABCD* mutants harbored the *flaA* expression plasmid (*P*_*lac*_*-P*_*flaA*_*-flaA*). This is another indication that mutants with increased levels of FlaA alone can drive motility to a limited extent but that efficient motility required a secondary flagellin. In *S. meliloti*, a feedback regulation has been reported to exist on flagellin synthesis, by which Δ*flaA* mutants have decreased expression of the secondary flagellins (12). We have not observed any striking effects on flagellar gene expression in the different *A. tumefaciens* flagellin mutants (Mohari and Fuqua, unpublished results).

Given that we were able to observe suppressors of severely compromised flagellin mutants with the Δ*flaA* deletion, it seemed likely that in the appropriate contexts secondary flagellins could compensate for the absence of FlaA. Suppression was observed only in mutants that retain a chromosomal copy of *flaB*, so we hypothesized that FlaA and FlaB might have somewhat similar functions. In the Δ*flaACD* fms-6 mutant, FlaBY129N in combination with base substitutions that alter the Shine-Dalgarno sequence reproduced the suppressor phenotype (Fig. 9B) but we discovered later that FlaBY129N is primarily responsible for rendering helical shape to flagellar filaments in these mutants (Fig. 10L). It is interesting that this kind of suppression was only observed in Δ*flaACD* suppressors where FlaB is the only flagellin present. However, for the Δ*flaACD* fms-1 mutant, FlaBY129N in combination with SciP G77R was required to reproduce the suppressor phenotype (Fig. 8). Perhaps similarly, the increased expression from the SD*-FlaB mutation in the fms-6 mutant in part contributed to the improved motility in Δ*flaACD*. The above scenario is a situation where the FlaBY129N mutation compensates for the absence of FlaA by rendering helical shape to the flagella and additional mutations can augment the impact of this change by increasing *flaB* expression.

Simple flagellar filaments of bacteria undergo polymorphisms that depend on the configuration of the protofilaments that make up the flagella. Key residues that facilitate these transformations have been documented (17). However, bacteria with complex filaments (e.g., *S. meliloti, A. tumefaciens*) are apparently not able to undergo polymorphisms of their flagellar filaments unless they are subjected to extremes of ionic strength and pH variations. Extreme pH values of 3 and 11 can transform helical filaments to straight ones (13). We found that mutation of a single tyrosine residue to an asparagine in FlaB (as in the fms-6 suppressor mutant) is sufficient to impart the helical shape of the mutant flagellar filament in our fms-6 suppressor mutant. Also, FlaA has an asparagine at the exact same position. Accordingly, we found that creating a FlaAN129Y (mutated) flagellin abolishes the helical flagellar conformation transforming them to straight filaments that do not function in the context of a Δ*flaACD* mutant (Figs. 9A, 10O-P). Hence it seems that FlaAN129 is a major contributor to the helical conformation to the flagella. It has been shown earlier in *Shewanella oneidensis* in which two flagellins FlaA and FlaB comprise the filaments, that two key residues determine the activity of the FlaB flagellin, however it is not known whether these residues determine helical shape (37). Here, we show that FlaAN129 is a critical residue that might contribute significantly to the role of FlaA as the major flagellin.

We observed that another frequent mechanism for suppression was via antagonism of SciP, a predicted inhibitor of flagellar gene expression. We have isolated three *A. tumefaciens* fms suppressors that are mutated in either the *sciP* (Atu2430) coding sequence or the predicted *sciP* Shine Dalgarno. The Δ*flaACD* fms-1 and Δ*flaAD* fms-12 suppressor mutants were found to be mutated in the helix turn helix (H-T-H) motif of SciP with L61F and G77R mutations respectively. The Δ*flaAC* fms-5 suppressor has a base substitution 9 bp upstream of *sciP* translational start site, in the Shine Dalgarno region (Fig. 7). *C. crescentus* SciP (small CtrA inhibitor protein) is an essential HTH transcriptional factor that directly binds to CtrA (27), a two-component signaling protein and the master regulator of flagellar gene control. As a Class I flagellar gene at the top of the flagellar gene hierarchy, CtrA regulates the Class II flagellar structural genes (28, 38). The SciP homologue from *A. tumefaciens* is 91% similar and 84% identical to that from *C. crescentus* and aligning them reveals that the helix turn helix motif is well conserved (Fig. S5). SciP exerts its effects in swarmer cells of *C. crescentus* and thus controls expression of many flagellar and chemotaxis genes by inhibiting CtrA, a function important for normal cell cycle progression. Several conserved residues including R35 and R40 outside of the HTH are critical for SciP-CtrA interaction and mutating these residues blocks the interaction thus hampering the SciP overexpression phenotypes (27). The residues mutated in the *A. tumefaciens* fms suppressors we isolated, have not been previously identified as important, however these are conserved and are clearly within the HTH, suggesting a role in DNA binding or for interactions with CtrA. The *sciP* homologue is not essential in *A. tumefaciens* and thus its role in regulating CtrA, which is itself essential, remains speculative (39). Even so, it is possible that the mutations are relieving the inhibition of SciP on CtrA to a certain extent, thus allowing greater transcription of flagellin genes analogously to the Shine-Dalgarno mutations, thus enhancing motility in the suppressors. The mutation immediately upstream of the *sciP* coding sequence changes the presumptive Shine-Dalgarno from (-11)TAGAGT(-6) to TATAGT, which is likely to decrease translational initiation, thereby diminishing SciP inhibition of flagellar gene expression.

Thus, there seem to be various mechanisms to achieve suppression and thus overcome the absence of specific flagellins. An open question is whether flagellar filaments are comprised of all four flagellins or whether they occur in specific combinations. There may be environmental conditions that lead to different proportions of the flagellins in filaments. We now know that the flagellins which comprise the flagellar filament of *A. tumefaciens* are not simply redundant in function, and we have gained insights into their specific roles and relative abundances.

## Materials and Methods

### Bacterial strains, plasmids and growth conditions

Table S2 lists all bacterial strains and plasmids used in this study. Table S3 lists all oligonucleotides. Buffers, reagents, antibiotics and media were obtained from Fisher Scientific (Pittsburgh, PA) and Sigma Chemical Co. (St. Louis, MO). Oligonucleotides were ordered from Integrated DNA Technologies (Coralville, IA). E.Z.N.A. Plasmid Miniprep kits (Omega Bio-tek, Norcross, GA) were used to purify DNA. All restriction enzymes and molecular biology reagents were obtained from New England Biolabs (Ipswich, MA). DNA was manipulated using standard techniques as described earlier (40). Standard Sanger-type DNA sequencing was performed on an ABI 3730 sequencer at the Indiana Molecular Biology Institute, Bloomington, IN. Plasmids were introduced into *A. tumefaciens* via electroporation (41, 42). *E. coli* was grown in LB medium and *A. tumefaciens* was grown in AT minimal medium (43) supplemented with 0.5% (w/v) glucose and 15 mM ammonium sulfate (ATGN). For allelic replacements, where ATSN medium was used, 5% (w/v) sucrose was used as the sole carbon source instead of glucose for *sacB* counterselection. The antibiotic concentrations used for *E. coli* were as follows in μg ml^−1^. 100 ampicillin (Amp) and 25 kanamycin (Km); for *A. tumefaciens* in μg ml^−1^: 300 kanamycin (Km), 50 spectinomycin (Sp). Isopropyl-thio-β-galactoside (IPTG) was added to the media for *P*_*lac*_ induction at the indicated concentrations, when appropriate.

### Construction of complementation constructs

Wild type or site-specifically mutated flagellin coding sequences (wild type *flaA, flaB, flaC* and *flaD* and their corresponding mutated genes: *flaA*T213C, *flaB*T190C, *flaC*T216C *and flaD*T276C) were amplified with primers comp P1 and comp P2 from AtC58 genomic DNA using Phusion High Fidelity DNA polymerase. These fragments were gel purified and ligated to pGEM-T Easy, their sequences were confirmed using appropriate restriction enzymes, then re-ligated to pSRK-Km previously cleaved with corresponding restriction enzymes. For all complementation constructs except the ones with the native flagellin promoters, 5’ primers were designed to incorporate the *NdeI* site of pSRK-Km in frame with the *lacZα* start codon to be cloned into the IPTG inducible expression vector pSRK-Km (44). The pSRK-Km constructs were electroporated into *A. tumefaciens* as described previously (41). For the construction of a *P*_*lac*_-*P*_*flaA*_*-flaA* and *P*_*lac*_*-P*_*flaA-*_*flaA*T213C the entire promoter region upstream of *flaA* along with its coding region was amplified using *flaA P*_*flaA*_ promoter comp P1 and flaA comp P2 primers. The 5’ *flaA* promoter comp P1 was designed with a stop codon TGA after the restriction site SpeI in order to terminate translation from *lacZ*α to prevent occlusion of the native translation start site. *P*_*lac*_-*P*_*native*_*-flaB* was similarly constructed.

### Construction of non-polar deletion mutants

Deletion mutants were constructed as described earlier (45). Primers for deleting the desired gene were carefully designed so that adjacent genes are not affected and to maintain potential translational coupling. DNA fragments were generated about 500-1000 bp upstream and downstream flanking the desired gene via PCR using P1 and P2 primers (for the upstream sequence) and with P3 and P4 primers (for the downstream sequence). Primers P2 and P3 were designed so that they could overlap with each other via 18 bp of complementarity (Table S3). The two flanking fragments were then joined together via a <u>s</u>plicing by <u>o</u>verlapping <u>e</u>xtension (SOEing) PCR reaction using Phusion high fidelity DNA polymerase (NEB, Ipswich, MA), as described earlier (46-48). All single deletion mutants were generated from WT C58 strain. For generation of Δ*flaAB* mutant, the Δ*flaA* mutant was used as the starting strain instead of the wild type C58 in order to ensure that Atu0544 ORF is retained in the corresponding mutants (Fig. S2). FlaAP1 and FlaBP2 were used as the P1 and P2 primers whereas FlaBP3 and FlaBP4 were used as the P3 and P4 primers. For subsequent generation of Δ*flaABC and* Δ*flaABD* mutants, the Δ*flaAB* mutant was used as the starting strain. The Δ*flaABC* mutant was generated with the use of FlaAP1 and FlaCP2 as P1 and P2 primers and with FlaCP3 and FlaCP4 as P3 and P4 primers. All multiple flagellin deletion mutants were generated starting from a single or double gene deletion mutant. The flanking DNA fragments were gel purified and another short PCR reaction of 5 cycles was performed using this as the template and primer. In order to join the fragments, the above product was amplified via another amplification using primers P1 and P4. This final product of ≈1 kb was ligated to pGEM-T Easy (Promega, Madison, WI). Colonies were confirmed by DNA sequencing, digested with the appropriate restriction enzymes and ligated into the suicide vector pNPTS138 cleaved with the same set of enzymes. The pNPTS138 plasmid ((45); MRK Alley, unpublished) has the Km resistance gene (Km^R^ and the *sacB* gene imparts sucrose sensitivity (Suc^S^). Since it has a *colE1* replication origin that does not function in *A. tumefaciens*, in order to form viable Km^R^ colonies, the plasmid must recombine into a stable endogenous replicon. Correct pNPTS138 derivatives were confirmed by digestion and the miniprep was transformed into the *E. coli* mating strain S17-λpir. The mating strain harboring the deletion plasmid was conjugated with *A. tumefaciens* via mating on LB, and recombinant colonies were selected for growth on ATGN Km plates. In order to confirm that the desired deletion plasmid has integrated and the colonies are sucrose sensitive, Km^R^ colonies were re-streaked on ATGN Km plates and ATSN Km plates. Colonies with normal growth on ATGN Km plates and very poor growth on ATSN Km plates were chosen for further processing. The next step was to select for the correct mutants that have lost the integrated plasmid due to a second recombination event and are sucrose resistant. To obtain these colonies, a single Km^R^/Suc^S^ colony was grown overnight in ATGN broth without Km selection and then plated on ATSN. The resultant Suc^R^ colonies were patched on ATSN and ATGN Km to verify plasmid excision. Diagnostic colony PCR was performed using primers P1 and P4 that flank the desired gene, to confirm that the target gene was deleted, and this amplification product was sequenced to confirm the deletion junction.

### Allelic replacement of wild type genes with the fms alleles

Allelic replacement procedures were performed as described previously (49). Wild type genes were replaced with either the fms or *sciP* allele in the parent flagellin mutant. The fragment containing the desired mutation in the middle was PCR amplified using Primers P1 fms and P2 fms or using Primers P1 *sciP* and P2 *sciP* from fms mutants – fms-1, fms-5, fms-6, fms-12 and fms-13 using Phusion High Fidelity DNA polymerase. Positioning the desired mutation in the center of the fragment increased the probability of recombination events on either side. The PCR fragments were gel purified and ligated to pGEM-T Easy. The remainder of the process was identical to the construction of non-polar deletion mutants described above. This included incorporation of the pNPTS138::fms allele(s) in the parent mutant *A. tumefaciens* via mating. For allelic replacement of *sciP* alleles, the procedure was modified a bit due to apparent toxicity of the gene when expressed in *E. coli*. We therefore used a tri-parental mating approach using the helper plasmid pRK2013;; (50) or pRK2073 (51). After desired Km^S^/Suc^R^ colonies were obtained – either from the bi-parental or the tri-parental mating, colony PCR was performed to amplify the DNA which was later gel purified and the desired fms or *sciP* mutation was confirmed via DNA sequencing using internal sequencing primers. The procedure of allelic replacement was repeated sequentially when more than one mutation needed to be incorporated in a parent mutant.

### Site-directed mutagenesis of flagellins

Site-directed mutagenesis was performed as described earlier for FlaAT213C, FlaCT216C and FlaDT276C, using the QuikChange protocol obtained from Stratagene Corp (49). Long, self-complementary mutagenic primers F1mut and R1mut (Tm ∼ 68°C) were designed harboring the desired mutations individually in the middle for all flagellin genes (Table S3). Mutagenic primers and Phusion DNA polymerase were used to perform PCR on the following pGEM-T easy plasmids: pBM139 (containing *flaA*), pBM147 (containing *flaC*) or pBM128 (containing *flaD*) (Table S3). *Dpn*I digestion of the PCR reactions was performed in order to remove the parental wild type and hemi-methylated plasmid DNA. The newly synthesized, uniformly non-methylated mutated DNA should not be cleaved, and will transform into *E. coli*, followed by sequencing to confirm the mutations and then introduced into wild type by allelic replacement as described above. The mutated DNA fragment (either *flaA*T213C or *flaC*T216C or *flaD*T276C) replaced its counterpart in the wild type genome. Allelic replacement for FlaBT190C strain was accomplished via a SOEing PCR reaction. Long, self-complementary mutagenic primers FlaBT190CF1mut and FlaBT190CR1mut (Tm ∼ 68°C) were designed harboring the desired mutation for T190C (*flaB*) (Table S3). As described above, about 500-1000 bp of upstream and downstream DNA sequences flanking the desired gene were PCR amplified with FlaB comp P1 and FlaBT190CR1mut primers (for the upstream sequence) and with FlaB comp P2 and FlaBT190CF1mut primers (for the downstream sequence). Owing to their self-complementarity, the two fragments were annealed, gel purified and ligated to pGEM-T Easy and the remaining procedure was identical to the protocol described above.

### Genome re-sequencing of *fms* mutants

After streak purifying the *fms* suppressors on ATGN, a paired-end library was prepared from their total genomic DNA using the modified protocol of Lazinski and Camilli (http://genomics.med.tufts.edu/home/sampl eprep) as described earlier (48). NEB Quick Blunting Kit (New England Biolabs) was used to blunt-end about 20 μg of sheared genomic DNA followed by the addition of one deoxyadenosine to their 3’ ends using the Klenow fragment of DNA Polymerase I. The NEB Quick Ligation Kit (New England Biolabs) was then used to ligate the DNA preparation with an adapter mix consisting of primers OLJ131 and OLJ137. The library was then amplified by PCR using the primers OLJ139 and OLJ140. Sequencing was performed on an Illumina MiSeq platform at the Center for Genomics and Bioinformatics at Indiana University.

### Motility assays

Petri dishes (100 mm) containing 0.25% ATGN motility agar were used to evaluate swimming proficiency. Swimming phenotypes were determined by inoculating fresh colonies into the center of the swim plates with 20 ml Bacto agar (BD, Sparks MD) (52). Antibiotics and IPTG were added when necessary. The plates were incubated at 28°C for up to seven days and their swim ring diameters were measured daily.

### Transmission Electron Microscopy

Transmission electron microscopy (TEM) was performed as described previously (22, 49). 300-mesh, 3-mm copper grids (Electron Microscopy Sciences, Hatfield, PA) were coated with carbon-formvar films. The sample for TEM was *A. tumefaciens* cells grown in ATGN to an OD_600_ ≈1.0 and diluted 1:10, 10 μl of which was applied to the formvar-coated grids. After 5 minutes, each grid was dried with filter paper and then negatively stained with 2% uranyl acetate for another 5 minutes. The grids were again dried to remove excess stain and then examined with a JEOL JEM-1010 transmission electron microscope set to 80 kV in the Indiana University Electron Microscopy Center.

### Visualization of flagellar filaments with Alexafluor 488 maleimide stain

*A. tumefaciens* wild type or mutant cells were grown up overnight and then subcultured until they reach exponential phase. Alexafluor 488 maleimide dye was diluted to 25 μg/ml in ATGN minimal media. 1-2 μl of diluted dye was mixed with the same volume of cells. The mixture was then added on a slide and incubated for 5-10 minutes with a cover slip on it. The samples were then observed at 60X magnification with a Nikon 90i microscope equipped with a Photometrics Cascade cooled CCD camera (excitation filter= 480/40 nm, dichromatic mirror= 505 nm, absorption filter = 535/50 nm).

### Amino acid sequence alignment and structural analysis

The FASTA *A. tumefaciens* FlaA, FlaB, FlaC And FlaD amino acid sequences were imported from Pubmed Protein to ClustalW2 and aligned as per the default settings (53). The FASTA for SciP were aligned using Clustal Omega with default settings (54). Phyre structures of the *A. tumefaciens* flagellins were individually generated using the *Salmonella* Typhimurium FliC flagellin structural template (PDB 3A5X). This structural analysis was important to create the site specific mutations in the flagellins (see Figs. S1 and S3A for details) (55).

### Mass Spectrometry

Flagellar filament preparation was done as described previously for *Caulobacter crescentus* (11). *A. tumefaciens* wild type and Δ*flaABC* mutant cells were grown overnight and then subcultured into 500 ml of ATGN minimal media and incubated till exponential phase. Cultures were then centrifuged at a speed of 10,000 x g for 15 min to remove cells. A fixed angle Beckman 70 Ti rotor was used to ultracentrifuge the resultant supernatant at 74,800 x g for 30 min. 20 ml of phosphate buffered saline (PBS) was used to suspend the pellet and the mixture was again centrifuged at 17,500 x g to remove any remaining cells. The supernatant was again ultracentrifuged at 74,800 x g for 30 min, to concentrate the preparation. 100 μl of PBS was used to suspend the pellet and the final filament preparation was examined for the presence of filaments via TEM.

Flagellar samples prepared as described above were analyzed via LC-MS/MS in the IU Laboratory for Biological Mass Spectrometry. These preparations were incubated for 45 min at 57 °C with 2.1 mM dithiothreitol to reduce cysteine residue side chains. These side chains were then alkylated with 4.2 mM iodoacetamide for 1 h in the dark at 21 °C. A solution containing 1 μg trypsin, in 25 mM ammonium bicarbonate was added and the samples were digested for 12 hours at 37 °C. The resulting peptides were desalted using a ZipTip (Millipore, Billerica MA) The samples were dried down and injected into an Eksigent HPLC coupled to a LTQ Orbitrap XL mass spectrometer (Thermo Fisher Scientific, Waltham MA) operating in “top five” data dependent MS/MS selection. The peptides were separated using a 75-micron, 15 cm column packed in-house with C18 resin (Michrom Bioresources, Auburn CA) at a flow rate of 300 nl/min. A one-hour gradient was run from Buffer A (2% acetonitrile, 0.1% formic acid) to 60% Buffer B (100% acetonitrile, 0.1% formic acid). Peptides shared between flagellin proteins as well as those specific to individual proteins were identified by MS database interpretation. For each peptide, LC peak areas were manually extracted using a +/− 0.1 Da mass window around the theoretical m/z value for the observed charge state. To determine the relative intensity of each flagellin protein to that of FlaB, MS -identified peptide pairs showing high sequence identity were compared. The ratio of the signal area for the peptide relative to its FlaB counterpart was determined. In the case of peptides containing a methionine residue, both the unmodified and oxidized version of the peptide were observed, and in these cases, the total signal area for both versions were calculated. One-way Anova analysis was used to calculate if there are any significant differences between the ratios of peptides.

### β-Galactosidase assays and *lacZ* fusions

The *lacZ* reporter gene was fused to the start codons of flagellin genes as described previously (56). Primers for *lacZ* fusion were designed so that the fragments had the ribosome-binding site and the gene start codon, in frame with the promoterless *lacZ* gene. Primers prom-1 and 2 (Table S3) were used to amplify promoter regions of *flaB* from *A. tumefaciens* wild type and fms-6 genomic DNA with Phusion DNA polymerase, followed by gel purification and ligation to pGEM-T Easy. Their sequences were verified, followed by restriction digestion that cleaved them at the designed restriction sites, to be ligated to the vector pRA301, cleaved with appropriate restriction enzymes. Following another round of sequence verification, the constructs were electroporated into *A. tumefaciens* Δ*flaACD* mutants as described earlier (41). β-galactosidase assays were performed to determine Miller units of at least three biological replicates as described previously (57).

## Acknowledgements

This project was supported by National Institutes of Health (NIH) grants GM080546 and GM120337 (C.F.). We thank Loubna Hammad for her contributions to the mass spectrometric analysis of flagellin preparations.

## REFERENCES

1. Chevance FF, Hughes KT. 2008. Coordinating assembly of a bacterial macromolecular machine. Nat Rev Microbiol 6:455–65.

2. Blair DF. 2003. Flagellar movement driven by proton translocation. FEBS Lett 545:86–95.

3. Turner L, Zhang R, Darnton NC, Berg HC. 2010. Visualization of flagella during bacterial swarming. J Bacteriol 192:3259–67.

4. Attmannspacher U, Scharf B, Schmitt R. 2005. Control of speed modulation (chemokinesis) in the unidirectional rotary motor of *Sinorhizobium meliloti*. Mol. Microbiol. 56:708–18.

5. Götz R, Schmitt R. 1987. *Rhizobium meliloti* swims by unidirectional, intermittent rotation of right-handed flagellar helices. J. Bacteriol. 169:3146–3150.

6. Götz R, Limmer N, Ober K, Schmitt R. 1982. Motility and chemotaxis in 2 strains of *Rhizobium* with complex flagella. J. Gen. Microbiol. 128:789–798.

7. Berg HC. 2003. The rotary motor of bacterial flagella. Annu. Rev. Biochem. 72:19–54.

8. Brown MT, Steel BC, Silvestrin C, Wilkinson DA, Delalez NJ, Lumb CN, Obara B, Armitage JP, Berry RM. 2012. Flagellar hook flexibility is essential for bundle formation in swimming *Escherichia coli* cells. J. Bacteriol. 194:3495–501.

9. Wilkinson DA, Chacko SJ, Venien-Bryan C, Wadhams GH, Armitage JP. 2011. Regulation of flagellum number by FliA and FlgM and role in biofilm formation by Rhodobacter sphaeroides. J Bacteriol 193:4010–4.

10. Kato S, Okamoto M, Asakura S. 1984. Polymorphic transition of the flagellar polyhook from *Escherichia coli* and *Salmonella typhimurium*. J. Mol. Biol. 173:463–476.

11. Faulds-Pain A, Birchall C, Aldridge C, Smith WD, Grimaldi G, Nakamura S, Miyata T, Gray J, Li G, Tang J, Namba K, Minamino T, Aldridge PD. 2011. Flagellin redundancy in *Caulobacter crescentus* and its implications for flagellar filament assembly. J. Bacteriol. doi:10.1128/jb.01172-10.

12. Scharf B, Schuster-Wolff-Buhring H, Rachel R, Schmitt R. 2001. Mutational analysis of the *Rhizobium lupini* H13-3 and *Sinorhizobium meliloti* flagellin genes: importance of flagellin A for flagellar filament structure and transcriptional regulation. J Bacteriol 183:5334–5342.

13. Scharf B. 2002. Real-time imaging of fluorescent flagellar filaments of *Rhizobium lupini* H13-3: flagellar rotation and pH-induced polymorphic transitions. J. Bacteriol. 184:5979–5986.

14. Tambalo DD, Bustard DE, Del Bel KL, Koval SF, Khan MF, Hynes MF. 2010. Characterization and functional analysis of seven flagellin genes in *Rhizobium leguminosarum* bv. *viciae*. Characterization of *R. leguminosarum* flagellins. BMC Microbiol 10:219.

15. Yen JY, Broadway KM, Scharf BE. 2012. Minimum requirements of flagellation and motility for infection of *Agrobacterium* sp strain H13-3 by flagellotropic bacteriophage 7-7-1. Appl. Environ. Microbiol. 78:7216–7222.

16. Chesnokova O, Coutinho JB, Khan IH, Mikhail MS, Kado CI. 1997. Characterization of flagella genes of *Agrobacterium tumefaciens*, and the effect of a bald strain on virulence. Mol. Microbiol. 23:579–590.

17. Hayashi F, Tomaru H, Furukawa E, Ikeda K, Fukano H, Oosawa K. 2013. Key amino acid residues involved in the transitions of L-to R-type protofilaments of the *Salmonella* flagellar filament. J. Bacteriol. 195:3503–3513.

18. Klose KE, Mekalanos JJ. 1998. Differential regulation of multiple flagellins in *Vibrio cholerae*. J. Bacteriol. 180:303–316.

19. Li C, Sal M, Marko M, Charon NW. 2010. Differential regulation of the multiple flagellins in Spirochetes. J Bacteriol 192:2596–603.

20. Deakin WJ, Parker VE, Wright EL, Ashcroft KJ, Loake GJ, Shaw CH. 1999. *Agrobacterium tumefaciens* possesses a fourth flagellin gene located in a large gene cluster concerned with flagellar structure, assembly and motility. Microbiology 145:1397–1407.

21. Blair KM, Turner L, Winkelman JT, Berg HC, Kearns DB. 2008. A molecular clutch disables flagella in the *Bacillus subtilis* biofilm. Science 320:1636–8.

22. Wang Y, Haitjema CH, and Fuqua C. 2014. The Ctp type IVb pilus locus of *Agrobacterium tumefaciens* directs formation of the common pili and contributes to reversible surface attachment. J. Bacteriol. 196:2979–2988.

23. Llewellyn M, Dutton RJ, Easter J, O’Donnol D, Gober JW. 2005. The conserved *flaF* gene has a critical role in coupling flagellin translation and assembly in *Caulobacter crescentus*. Mol Microbiol 57:1127–42.

24. Anderson PE, Gober JW. 2000. FlbT, the post-transcriptional regulator of flagellin synthesis in *Caulobacter crescentus*, interacts with the 5 ’ untranslated region of flagellin mRNA. Mol. Microbiol. 38:41–52.

25. Ferooz J, Lemaire J, Letesson JJ. 2011. Role of FlbT in flagellin production in *Brucella melitensis*. Microbiology 157:1253–62.

26. Mangan EK, Malakooti J, Caballero A, Anderson P, Ely B, Gober JW. 1999. FlbT couples flagellum assembly to gene expression in *Caulobacter crescentus*. J. Bacteriol. 181:6160–6170.

27. Gora KG, Tsokos CG, Chen YE, Srinivasan BS, Perchuk BS, Laub MT. 2010. A cell-type-specific protein-protein interaction modulates transcriptional activity of a master regulator in *Caulobacter crescentus*. Mol. Cell 39:455–467.

28. Tan MH, Kozdon JB, Shen X, Shapiro L, McAdams HH. 2010. An essential transcription factor, SciP, enhances robustness of *Caulobacter* cell cycle regulation. Proc. Natl. Acad. Sci. USA 107:18985–18990.

29. Kuwajima G, Asaka JI, Fujiwara T, Fujiwara T, Node K, Kondo E. 1986. Nucleotide-sequence of the *hag* gene encoding flagellin of *Escherichia coli*. J. Bacteriol. 168:1479–1483.

30. Enomoto M, Sakai A, Tominaga A. 1985. Expression of an *Escherichia coli* flagellin gene, *hag48*, in the presence of a *Salmonella* H1-repressor. Mol. Gen. Genet. 201:133–135.

31. Cohen-Krausz S, Trachtenberg S. 1998. Helical perturbations of the flagellar filament: *Rhizobium lupini* H13-3 at 13 angstrom resolution. J. Struct. Biol. 122:267–282.

32. Felix G, Duran JD, Volko S, Boller T. 1999. Plants have a sensitive perception system for the most conserved domain of bacterial flagellin. Plant J 18:265–76.

33. Zipfel C, Kunze G, Chinchilla D, Caniard A, Jones JD, Boller T, Felix G. 2006. Perception of the bacterial PAMP EF-Tu by the receptor EFR restricts Agrobacterium-mediated transformation. Cell 125:749–60.

34. Renault TT, Abraham AO, Bergmiller T, Paradis G, Rainville S, Charpentier E, Guet CC, Tu Y, Namba K, Keener JP, Minamino T, Erhardt M. 2017. Bacterial flagella grow through an injection-diffusion mechanism. Elife 6.

35. Iida Y, Hobley L, Lambert C, Fenton AK, Sockett RE, Aizawa S. 2009. Roles of multiple flagellins in flagellar formation and flagellar growth post bdelloplast lysis in *Bdellovibrio bacteriovorus*. J Mol Biol 394:1011–21.

36. Millikan DS, Ruby EG. 2004. *Vibrio fischeri* flagellin A is essential for normal motility and for symbiotic competence during initial squid light organ colonization. J Bacteriol 186:4315–25.

37. Sun L, Dong Y, Shi M, Jin M, Zhou Q, Luo Z-Q, Gao H. 2014. Two residues predominantly dictate functional difference in motility between *Shewanella oneidensis* flagellins FlaA and FlaB. J. Biol. Chem. 289:14547–14559.

38. Laub MT, Chen SL, Shapiro L, McAdams HH. 2002. Genes directly controlled by CtrA, a master regulator of the *Caulobacter* cell cycle. Proc. Natl. Acad. Sci. USA 99:4632–4637.

39. Curtis PD, Brun YV. 2014. Identification of essential alphaproteobacterial genes reveals operational variability in conserved developmental and cell cycle systems. Mol Microbiol 93:713–35.

40. Sambrook J, Fritsch EF, Maniatis T. 1989. Molecular cloning a laboratory manual second edition vols. 1 2 and 3.

41. Mersereau M, Pazour GJ, Das A. 1990. Efficient transformation of *Agrobacterium tumefaciens* by electroporation. Gene 90:149–151.

42. Fuqua WC, Winans SC. 1994. A LuxR-LuxI type regulatory system activates *Agrobacterium* Ti plasmid conjugal transfer in the presence of a plant tumor metabolite. J. Bacteriol. 176:2796–2806.

43. Tempe J, Petit A, Holsters M, Montagu MV, Schell J. 1977. Thermosensitive step associated with transfer of Ti plasmid during conjugation - possible relation to transformation in crown gall. Proc. Natl. Acad. Sci. USA 74:2848–2849.

44. Khan SR, Gaines J, Roop RM, II, Farrand SK. 2008. Broad-host-range expression vectors with tightly regulated promoters and their use to examine the influence of TraR and TraM expression on Ti plasmid quorum sensing. Appl. Environ. Microbiol. 74:5053–5062.

45. Morton ER, Fuqua C. 2012. Genetic manipulation of *Agrobacterium*. Curr Protoc Microbiol doi:10.1002/9780471729259.mc03d02s25:Unit 3D 2.

46. Warrens AN, Jones MD, Lechler RI. 1997. Splicing by overlap extension by PCR using asymmetric amplification: An improved technique for the generation of hybrid proteins of immunological interest. Gene 186:29–35.

47. Merritt PM, Danhorn T, Fuqua C. 2007. Motility and chemotaxis in *Agrobacterium tumefaciens* surface attachment and biofilm formation. J. Bacteriol. 189:8005–8014.

48. Kim J, Heindl JE, Fuqua C. 2013. Coordination of division and development influences complex multicellular behavior in *Agrobacterium tumefaciens*. PLoS One 8:doi:10.1371/journal.pone.0056682.

49. Mohari B, Licata NA, Kysela DT, Merritt PM, Mukhopadhyay S, Brun YV, Setayeshgar S, Fuqua C. 2015. Novel pseudotaxis mechanisms improve migration of straight-swimming bacterial mutants through a porous environment. mBio 6:e00005–15.

50. Figurski DH, Helinski DR. 1979. Replication of an origin-containing derivative of plasmid RK2 dependent on a plasmid function provided in trans. Proc. Natl. Acad. Sci. USA 76:1648–1652.

51. Ditta G. 1986. TN5 mapping of *Rhizobium* nitrogen-fixation genes. Methods Enzymol. 118:519–528.

52. Adler J. 1966. Chemotaxis in bacteria. Science 153:708–716.

53. Larkin MA, Blackshields G, Brown NP, Chenna R, McGettigan PA, McWilliam H, Valentin F, Wallace IM, Wilm A, Lopez R, Thompson JD, Gibson TJ, Higgins DG. 2007. Clustal W and clustal X version 2.0. Bioinformatics 23:2947–2948.

54. Sievers F, Wilm A, Dineen D, Gibson TJ, Karplus K, Li W, Lopez R, McWilliam H, Remmert M, Söding J, Thompson JD and Higgins DG. 2011. Fast, scalable generation of high-quality protein multiple sequence alignments using Clustal Omega. Mol Syst Biol 7:539.

55. Kelley LA, Mezulis S, Yates CM, Wass MN, Sternberg MJE. 2015. The Phyre2 web portal for protein modeling, prediction and analysis. Nat Protocol 10:845–858.

56. Xu J, Kim J, Koestler BJ, Choi J-H, Waters CM, Fuqua C. 2013. Genetic analysis of *Agrobacterium tumefaciens* unipolar polysaccharide production reveals complex integrated control of the motile-to-sessile switch. Mol. Microbiol. 89:929–948.

57. Heckel BC, Tomlinson AD, Morton ER, Choi JH, Fuqua C. 2014. *Agrobacterium tumefaciens exoR* controls acid response genes and impacts exopolysaccharide synthesis, horizontal gene transfer, and virulence gene expression. J Bacteriol 196:3221–33.

